# METTL3-mediated m^6^A modification of DNMT1 enhances ovarian cancer progression

**DOI:** 10.1101/2025.09.10.675425

**Authors:** Kanchan Kumari, Iván Pérez-Núñez, Aleix Noguera, Raquel Pluvinet, Carlos Peula, Annika Patthey, Ghazala Naz Khan, Kinga Pajdzik, Ángel-Carlos Román, Erik Dassi, Chuan He, Eva Lundin, Manel Esteller, Toni Celià-Terrassa, Francesca Aguilo

## Abstract

High-grade serous carcinoma (HGSC), the most lethal subtype of ovarian cancer, is often diagnosed at advanced stages owing to its asymptomatic progression and lack of early detection markers. In this study, we identified a critical oncogenic role of the RNA methyltransferase METTL3 and the N^6^-methyladenosine (m^6^A) RNA modification pathway in HGSC. Depletion of METTL3 or its binding partner METTL14 impairs ovarian cancer proliferation and tumor progression. Mechanistically, m^6^A deposition enhances the translation of DNA methyltransferase DNMT1, an epigenetic repressor that silences tumor suppressor genes. Pharmacologic inhibition of DNMT1 led to DNA hypomethylation and upregulation of the tumor suppressors TNFAIP3 and FBXO32. Consistently, METTL3 depletion also increased the expression of these genes supporting a model in which METTL3 sustains oncogenesis by maintaining DNMT1 protein levels and repressing anti-tumor pathways. These findings position METTL3-mediated RNA modifications and DNMT1 as promising therapeutic targets in HGSC.

## Introduction

Ovarian cancer is the most lethal gynaecological malignancy and ranks as the fifth leading cause of cancer-related deaths among women [1]. Ovarian carcinomas are classified into various histological subtypes based on their origin [2]. High-grade serous carcinoma (HGSC) is the most common and aggressive subtype, often diagnosed at advanced stages with widespread metastasis [3], highlighting the need to elucidate the molecular mechanisms driving HGSC progression.

RNA modifications, including N6-methyladenosine (m^6^A), are critical regulators of gene expression, and their dysregulation contributes to cancer development and progression [4, 5]. m^6^A is catalyzed by a methyltransferase complex composed of METTL3, the catalytic core, and METTL14, along with other proteins such as WTAP, RBM15, VIRMA, and CBLL1, collectively referred to as “writers”. m^6^A is a reversible mark that can be removed by demethylases, known as “erasers”, such as fat mass and FTO and ALKBH5 [6], and recognized by “readers” including YTHDF1/2/3, YTHDC1/2, IGF2BP1/2/3, and HNRNPA2B1, which direct the fate of modified transcripts [7].

Among these RNA-modifying proteins, METTL3 has emerged as a key oncogene in ovarian cancer. Elevated METTL3 expression has been consistently observed in ovarian cancer tissues and cells [8, 9]. Functional studies have demonstrated that METTL3 promotes oncogenesis by activating pathways such as PI3K/AKT/mTOR via miR-126-5p-mediated PTEN inhibition [8], enhancing AXL translation, and stabilizing oncogenic RNAs including *MALAT1*, *DDR2*, and *RHPN1-AS1* via m^6^A modifications [9–12]. Additionally, METTL3 targets miR-1246 to suppress CCNG2 expression, thereby supporting tumor metabolism and proliferation [13]. However, the role of METTL14 in ovarian cancer remains ambiguous, with conflicting studies of tumor-promoting and suppressive effects [14, 15].

In this study, we explored the roles of METTL3 and METTL14 in HGSC using an OVCAR8 cell line model. Knockdown of either protein significantly impaired tumor cell progression, and transcriptomic profiling revealed widespread deregulation of cancer related pathways. m^6^A immunoprecipitation followed by sequencing (MeRIP sequencing) identified DNMT1—a key transcriptional repressor—as an m^6^A-modified target. Despite the unchanged transcripts levels, METTL3 depletion reduced DNMT1 protein levels through impaired translation. Functional assays further showed that pharmacological inhibition of DNMT1 using GSK-3484862 upregulated the tumor suppressors TNFAIP3 and FBXO32-mirroring the effects of METTL3 knockdown. Together, these findings highlight a METTL3-DNMT1 regulatory axis that facilitates ovarian cancer progression by enhancing DNMT1 translation. Targeting this pathway maybe a promising therapeutic strategy for HGSC treatment.

## Results

### METTL3 and METTL14 drive ovarian tumorigenesis

To investigate the role of METTL3 and METTL14 in ovarian cancer, we first performed knockdown experiments in the HGSC OVCAR8 cell line using short-hairpin RNAs (shRNAs) (referred to as sh1 and sh2 for each gene) along with a scrambled control (Scr). Knockdown efficiency was validated by western blot and reverse transcription quantitative PCR (RT-qPCR) (**Fig. S1A-D**). The depletion of either METTL3 or METTL14 significantly reduced cell proliferation (**Fig. 1A,B**), colony formation (**Fig. 1C-E**) and migration (**Fig. 1F-H**). Annexin V staining revealed increased apoptosis (**Fig. 1I,J**), whereas cell cycle analysis showed G0/G1 arrest and reduced G2/M populations (**Fig. 1K,L**).

**Figure 1.**
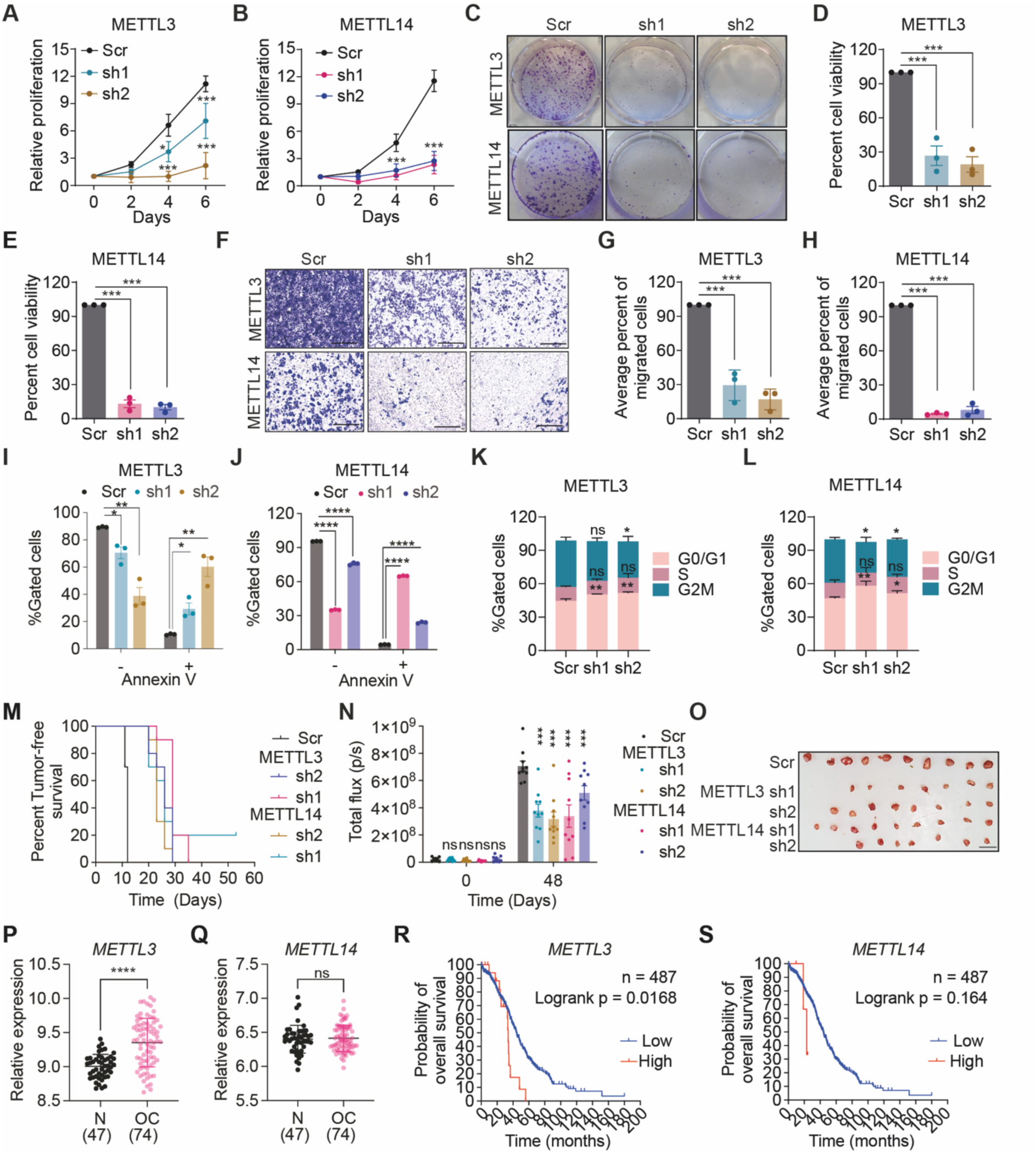
METTL3 and METTL14 promote ovarian cancer cell growth, migration, and poor patient prognosis. **A, B**. Cell proliferation rates of METTL3-(**A**) and METTL14-depleted (**B**) cells measured over 6 days. **C**. Representative image of a colony formation assay with METTL3- and METTL14-depleted cells, evaluated over 10 days. **D, E**. Bar graphs illustrating relative cell viability, calculated from absorbance values of dissolved crystal violet stain at 570 nm, corresponding to the colony formation assay in panel (**C**). **F**. Representative trans-well assay image showing the migration of METTL3- and METTL14-depleted cells. Scale bar, 200 µm. **G, H**. Quantification of migrated cells from panel (**F**) using ImageJ analysis. **I, J**. Graphs depicting the proportion of live (Annexin V-) and apoptotic (Annexin V+) cells in METTL3- (**I**) and METTL14-depleted (**J**) cells relative to Scr. **K, L**. Cell cycle distribution of cells following METTL3 (**K**) and METTL14 knockdown (**L**). **M**. Kaplan-Meier survival curve representing tumor-free survival of mice subcutaneously transplanted with Scr or METTL3- and METTL14-depleted OVCAR8 cells. **N**. Quantification of bioluminescence signal over the tumor region at day 0 and day 48, indicating tumor progression in each group. **O**. Representative images of tumors collected at the end of the experiment, with scale bar = 1 cm. **P**. Scatter plots representing *METTL3* (**P**) and *METTL14* (**Q**) mRNA levels in normal and ovarian cancer tissues (GEO dataset GSE66957). **R, S**. Kaplan-Meier survival curves for overall survival for METTL3 (**R**) and METTL14 (**S**) expression in high-grade serous carcinoma (HGSC) using TCGA data. Data are presented as mean ± SD or representative images of n > 3 independent biological experiments. 10 tumors per experimental group were included in the study. Data are presented as mean ± SD or as representative images from more than three independent biological experiments. Statistical significance was determined using Student’s t-test for comparisons between two groups, and one-way ANOVA for comparisons among three or more groups. ns = not significant (p > 0.05); *p < 0.05; **p < 0.01; ***p < 0.001; ****p< 0.0001.

Pharmacological inhibition of METTL3 using STM2457 elicited comparable, albeit milder, effects consistent with the notion that METTL3 can also promote tumorigenesis through m^6^A-independent mechanisms that are independent of m^6^A methyltransferase activity [16, 17]. STM2457 treatment significantly impaired OVCAR8 cell proliferation, colony formation, and migration (**Fig S1E-J**), while increasing apoptotic population of cells that were not statistically significant (**Fig. S1K**). Additionally, STM2457 altered cell cycle progression, reducing the proportion of cells in the G0/G1 and G2/M phases while increasing the S phase population (**Fig. S1L**). Notably, METTL3 silencing reduced METTL14 protein levels without affecting its mRNA levels, whereas METTL14 knockdown similarly decreased METTL3 protein abundance (**Fig. S1A,B; S2A-C**), suggesting mutual post-transcriptional stabilization, as previously observed in other cellular models [18]. Supporting this, METTL14 was detected by immunoprecipitation of endogenous METTL3 (**Fig. S2D**). To further investigate whether METTL3 stabilizes METTL14 in ovarian cancer, we generated doxycycline-inducible, shRNA-resistant, FLAG-tagged METTL3 in OVCAR8 cells. Reexpression of METTL3 restored the reduced METTL14 levels caused by METTL3 knockdown (**Fig. S2E**). Moreover, treatment of cells with the proteasome inhibitor MG132 increased METTL14 protein levels in METTL3 knockdown cells (**Fig. S2F,G**), indicating that METTL14 undergoes proteasomal degradation in METTL3-deficient cells, as previously suggested [18]. METTL3 depletion also lowered the protein levels of other writer complex components, such as WTAP and VIRMA, but not HAKAI, underscoring the role of METTL3 in preserving the integrity of the m^6^A methyltransferase complex (**Fig S2H**).

To assess the tumorigenic roles of METTL3 and METTL14 in vivo, luciferase-labelled OVCAR8 cells, with either knockdown of METTL3, METTL14 or scrambled control were subcutaneously injected into nude mice. Mice injected with control OVCAR8 cells rapidly developed tumors, with a median tumor-free survival of 12 days. In contrast, mice injected with METTL3- or METTL14-depleted cells exhibited significantly delayed tumor onset, with tumor-free survival extending to 21 and 28 days, respectively (**Fig. 1M**). Bioluminescence imaging further demonstrated that tumors from METTL3- and METTL14-deficient cells grew at a markedly reduced rate compared to control tumors (**Fig. 1N**). After eight weeks, excised tumors showed significantly lower volume and weight in the knockdown groups than in the controls (**Fig. 1O**; **S2I**,**J**), highlighting the critical role of METTL3 and METTL14 in promoting tumor growth and progression in ovarian cancer in vivo.

To assess METTL3 and METTL4 expression in ovarian cancer, we analyzed mRNA levels in 74 ovarian carcinomas and 47 normal ovarian samples from the GSE37582 dataset of the Gene Expression Omnibus (GEO). *METTL3* was significantly upregulated in ovarian cancer tissues compared to normal tissues whereas *METTL14* did not show a significant change (**Fig. 1P,Q**). Kaplan-Meier (KM) survival analyses using data from The Cancer Genome Atlas (TCGA) showed that high *METTL3* expression correlated with reduced overall survival across all ovarian cancer subtypes (**Fig. S2K**). Although elevated METTL14 expression was associated with better outcomes, this trend was not statistically significant (**Fig. S2L**). Specifically, in HGSC, we found that high *METTL3* levels predicted poorer survival (**Fig. 1R**), whereas high METTL14 expression showed a non-significant trend toward worse prognosis (**Fig. 1S**).

Collectively, these findings underscore the critical roles of METTL3 and METTL14 as key drivers of ovarian cancer cell growth and migration. The impact of STM2457 highlights the therapeutic potential of targeting METTL3, which is supported by its upregulation and association with poor prognosis. In contrast, despite its functional importance, METTL14 lacks clear prognostic relevance.

### Transcriptomic impact of METTL3 and METTL14 depletion and global mapping of m^6^A in OVCAR8 cells

To explore the transcriptional changes associated with METTL3 and METTL14 depletion, we performed RNA sequencing (RNA-seq) analysis on OVCAR8 cells transduced with control or shRNAs targeting METTL3 and METTL14. Following METTL3 silencing, we identified 3,448 and 4,896 differentially expressed genes in sh1 and sh2, respectively (**Fig. 2A, S3A,B; Table S1**). Among these, 743 were upregulated, and 705 were downregulated (**Fig. S3C,D**). METTL14 depletion resulted in 6,117 and 5,146 differentially expressed genes for sh1 and sh2, respectively (**Fig. 2B; S3E,F; Table S2**) with 905 genes upregulated and 994 downregulated across both shRNAs (**Fig. S3G,H**). Gene Ontology (GO) enrichment analysis revealed that the upregulated genes were involved in a broad range of biological processes, including regulation of transcription by RNA polymerase II, chromatin remodelling, protein transport, ubiquitination and in-utero embryonic development (**Fig. 2C**). Conversely, downregulated genes were more selectively enriched in cancer-associated pathways, such as signal transduction, regulation of cell proliferation, extracellular matrix organization, and negative regulation of cell migration (**Fig. 2D**). To determine whether these transcriptional changes are linked to m^6^A RNA methylation, we performed m^6^A RNA immunoprecipitation followed by sequencing (MeRIP-seq) in OVCAR8 cells to directly map m^6^A-modified transcripts in the HGSC context. A total of 6,178 m^6^A peaks were identified, primarily deposited near the stop codons of protein-coding transcripts and within the coding sequence (CDS) of the transcripts (**Fig. 2E,F; Table S3**). Motif analysis confirmed the presence of canonical GGAC consensus motif within the peaks, further validating the specificity of our MeRIP-seq data (**Fig. 2G**). GO was enriched in pathways critical to cancer, including transcriptional regulation, apoptotic processes, cell cycle control, DNA repair, cell migration, in-utero embryonic development, mRNA transport, and regulation of TGFβ receptor signalling (**Fig. 2H**), supporting a key role of m^6^A modification in modulating gene expression programs in ovarian cancer.

**Figure 2.**
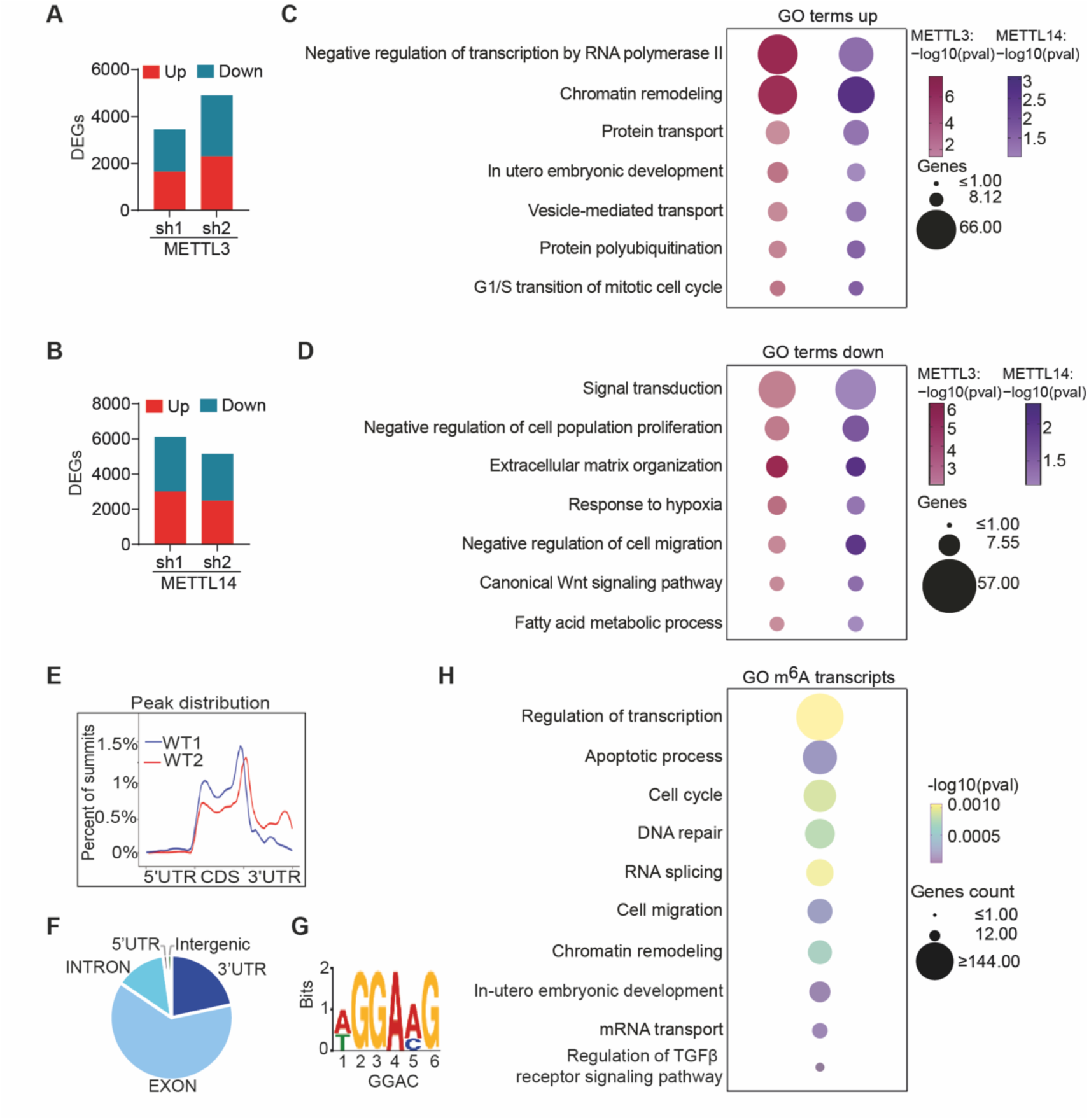
METTL3 and METTL14 regulate gene expression and m^6^A-modified transcripts involved in key biological processes in OVCAR8 Cells. **A, B.** Bar graph shows the comparison of differentially expressed genes between different shRNAs targeting METTL3 (**A**) and METTL14 (**B**). **C, D.** GO Biological process enrichment analysis of shared upregulated (**C**), and downregulated (**D**), genes after METTL3 and METTL14 depletion, visualized by dot plot. **E.** Density distribution of m^6^A peaks across the mRNA transcripts, including 5’ untranslated regions (5’UTR), coding sequence (CDS) and 3’UTR. **F**. Pie chart illustrating the distribution of overlapping m^6^A peaks across the mRNA transcriptome. **G**. m^6^A motif detected by the MEME motif analysis with MeRIP-seq data in OVCAR8 cells. **H**. GO dot plot showing enriched biological functions of m^6^A modified transcripts identified in MeRIP-seq data. Representative images of n=2 independent biological experiments.

### m^6^A promotes DNMT1 translation

MeRIP-seq analysis revealed that *DNMT1*, an essential enzyme that maintains genomic DNA methylation, is m^6^A-modified within its coding sequence (CDS), specifically at exon 8. (**Fig 3A**). Since DNA methylation is crucial for gene expression, chromatin stability, and genetic imprinting and may represent an early step in neoplastic transformation, we investigated the regulation of DNMT1 by m^6^A in ovarian cancer [19]. m^6^A RNA immunoprecipitation followed by qPCR showed a 2.5-fold enrichment of m^6^A-modified *DNMT1* mRNA (**Fig. 3B**), validating the MeRIP-seq results. For nuclepotide specific analysis, the m^6^A site within exon 8 at position 3013 was assessed using SELECT, a robust method for detecting locus-specific m^6^A modifications [20]. STM2457-treated OVCAR8 cells showed a 3-fold increase in SELECT products, indicating a reduced m^6^A modification at this site (**Fig 3C**). Similarly, METTL3 and METTL14-depleted OVCAR8 cells displayed enhanced SELECT products (**Fig. 3D, S4A**)

**Figure 3:**
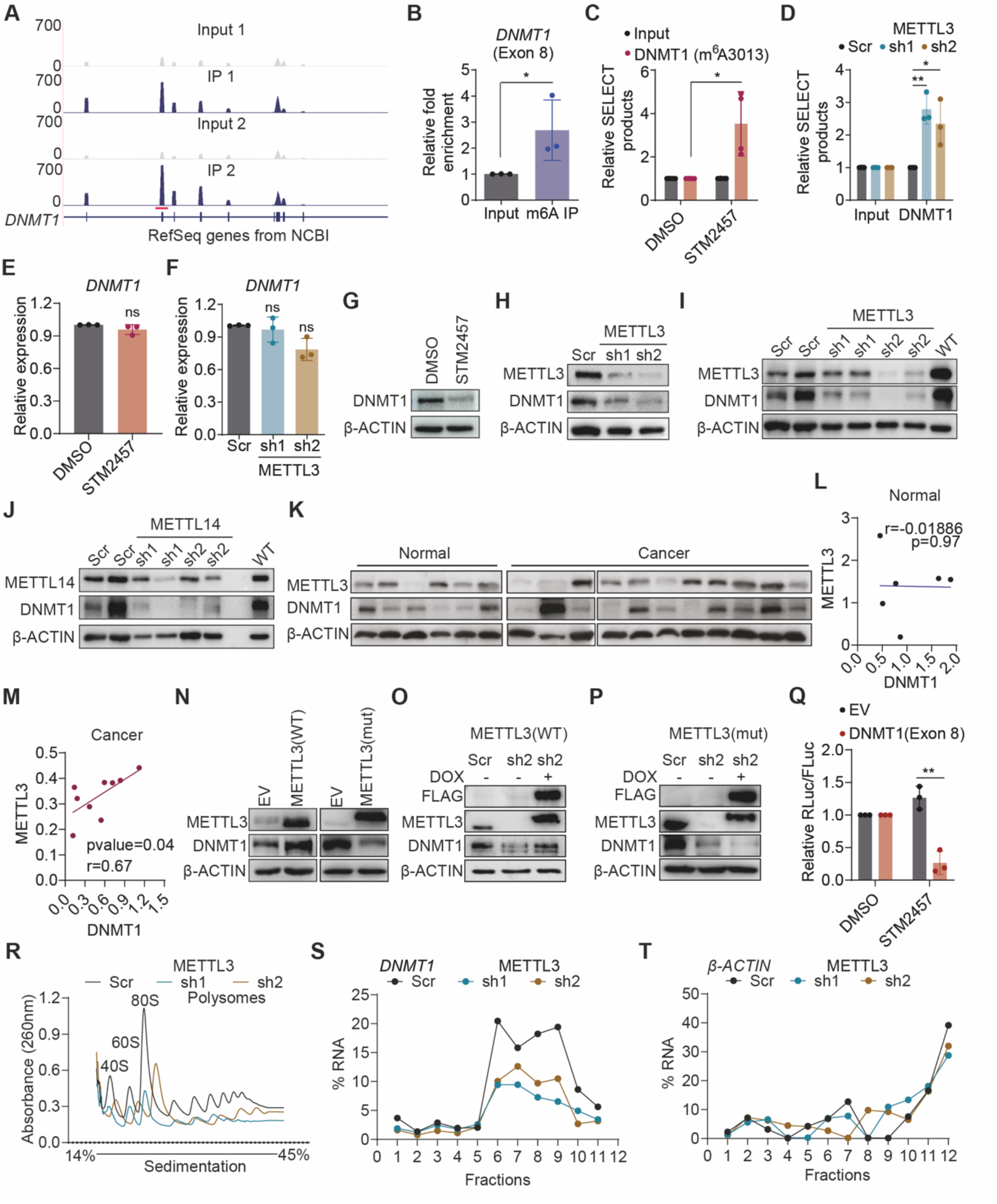
METTL3-mediated m^6^A modification promotes DNMT1 translation. **A**. Integrative Genomics Viewer (IGV) track demonstrating the distribution of m^6^A peaks across exon 8 of *DNMT1* mRNA, comparing m^6^A immunoprecipitated RNA (blue peaks) to the input control (gray peaks). **B**. Bar graph showing fold enrichment of DNMT1 following m^6^A IP compared to the input control. **C, D**. Bar graphs illustrating the levels of SELECT products after STM2457 treatment (**C**) and METTL3 depletion (**D**). **E, F.** qPCR analysis of *DNMT1* mRNA expression upon STM2457 treatment (**E**), and METTL3 depleted cells (**F**), visualized as bar graphs. **G, H.** Western blotting showing DNMT1 protein levels in STM2457 treated (**G**) and METTL3-depleted cells (**H**). **I**, **J.** Western blot analysis of DNMT1 expression in protein extracts from xenografts generated from METTL3- (**I**) and METTL14-depleted (**J**) cells. **K.** Expression of METTL3 and DNMT1 in normal ovarian epithelial and ovarian cancer tissues. **L**, **M**. Correlation plot of DNMT1 and METTL3 protein expression in protein extracts of normal ovarian epithelial patients (**L**) and ovarian cancer patient tissues (**M**) quantified from (**K**). **N**. Western blot images depict the expression level of DNMT1 upon overexpression of wild-type (WT) METTL3 and catalytic site mutated (mut) METTL3. **O**. Western blotting of FLAG, METTL3 and DNMT1 from whole-cell extracts from scr and METTL3-knockdown (sh2) cells and upon expression of wild-type cytoplasmic METTL3 (sh2 + doxycycline (dox)) cells. **P**. Western blot of FLAG, METTL3 and DNMT1 from whole-cell extracts from scr and METTL3-knockdown (sh2) cells, and upon expression of catalytically dead cytoplasmic METTL3 (sh2 + dox) cells. **Q**. Relative Renilla luciferase activity (RLuc) of psiCHECK2-DNMT1Exon 8 in cells treated with DMSO (control) or STM2457 for 48 h. RLuc was measured and normalized to firefly luciferase activity (FLuc). **R**. Polysome profiling curve for METTL3-depleted cells illustrating the global ribosomal distribution. **S, T.** qPCR analysis of *DNMT1* (**S**) and *β-actin* (**T**) mRNA in polysome fractions following profiling (**R**). For western blot analysis, β-ACTIN was used as the loading control. Data are presented as the mean ± SD or as representative images from more than three independent biological experiments. Statistical significance was determined using the Student’s t-test for comparisons between two groups, and one-way ANOVA for comparisons among three or more groups. ns = not significant (p > 0.05); *p < 0.05; **p < 0.01.

To investigate the fate of m^6^A-modified *DNMT1* mRNA, we first focused on its RNA stability. Following actinomycin D treatment, *DNMT1* mRNA levels remained stable over a 48-hour treatment period in both control and STM2457-treated cells as well as in METTL3-depleted cells (**Fig. S4B,C**), indicating that m^6^A did not influence *DNMT1* transcript stability. Next, we assessed DNMT1 protein levels after METTL3 and METTL14depletion as well as STM2457 treatment. A significant reduction in DNMT1 protein levels without corresponding changes in mRNA levels was observed (**Fig. 3E-H, S4D,E**). Notably, METTL14 depletion with shRNA_1 increased *DNMT1* mRNA levels (**Fig. S4D**), suggesting a potential feedback regulation. Moreover, DNMT1 protein expression was abrogated in the protein extracts from METTL3- and METTL14-depleted mouse tissues (**Fig. 3I,J**). A significant positive correlation between METTL3 and DNMT1 expression was also observed in ovarian cancer samples, but not in normal tissues, supporting a cancer-specific role for m^6^A methylation in regulating DNMT1 expression (**Fig. 3K-M**).

To assess the specificity of the observed phenotypes rescue experiments were performed. Overexpression of wild-type (WT) METTL3 resulted in a significant increase in DNMT1 protein levels, whereas overexpression of the catalytically inactive the METTL3 mutant showed reduced DNMT1 protein levels (**Fig. 3N**). Moreover, in OVCAR8 cells expressing FLAG-tagged, shRNA-resistant, doxycycline-inducible METTL3 cDNA, DNMT1 protein levels were restored following METTL3 knockdown (**Fig. 3O**). In contrast, the expression of a catalytically inactive METTL3 construct in the same context further reduced DNMT1 levels compared to that in cells with METTL3 knockdown alone (**Fig. 3P**). These findings confirm that the m^6^A activity of METTL3 is required to maintain DNMT1 protein expression. Consistently, METTL14 overexpression in OVCAR8 cells led to increased DNMT1 protein levels, further supporting the role of the m^6^A methyltransferase complex in sustaining DNMT1 expression (**Fig. S4F**).

To further investigate whether m^6^A modification regulates DNMT1 translation, we transfected OVCAR8 cells with a dual-luciferase reporter construct containing the *DNMT1* Exon 8 sequence downstream of the Renilla luciferase gene. In this construct, the consensus m^6^A motifs within both the Renilla and Firefly luciferase genes were ablated to eliminate potential confounding effects from endogenous methylation. Upon STM2457 treatment, Renilla luciferase activity was significantly reduced, suggesting that m^6^A modification within *DNMT1* Exon 8 was sufficient to influence its translational output (**Fig. 3Q**). To further support this hypothesis, we conducted polysome profiling of METTL3-depleted cells. METTL3 depletion led to a global reduction in ribosomal subunits, monosomes, and polysomes compared to the control cells (**Fig. 3R; S4G,H**), suggesting an overall impairment in translation. Subsequent qPCR analysis of polysome fractions revealed a marked decrease in *DNMT1* mRNA-associated polysomes (**Fig. 3S; S4I,J)**, whereas *β-ACTIN* mRNA, serving as a control, remained unaffected (**Fig. 3T; S4K,L)**. Collectively, these findings support a model in which m^6^A modification enhances DNMT1 translation, highlighting a key post-transcriptional mechanism by which METTL3 regulates DNMT1 protein levels.

### DNMT1 promotes ovarian cancer growth

Given that METTL3 promotes DNMT1 translation without affecting its mRNA expression, we next sought to determine the functional relevance of DNMT1 in ovarian cancer. We first assessed DNMT1 expression levels using the GEO dataset GSE37582, which revealed no significant difference in *DNMT1* mRNA levels between ovarian cancer samples and normal tissues (**Fig. 4A**). Similarly, analysis of TCGA datasets showed no statistically significant association between *DNMT1* expression and overall survival across all ovarian cancer subtypes (**Fig. 4B**). However, there was a non-significant trend indicating poorer survival with increased *DNMT1* expression (**Fig. 4C**), suggesting a potential involvement of DNMT1 in HGSC progression that might not be fully captured by transcript-level data alone.

**Figure 4.**
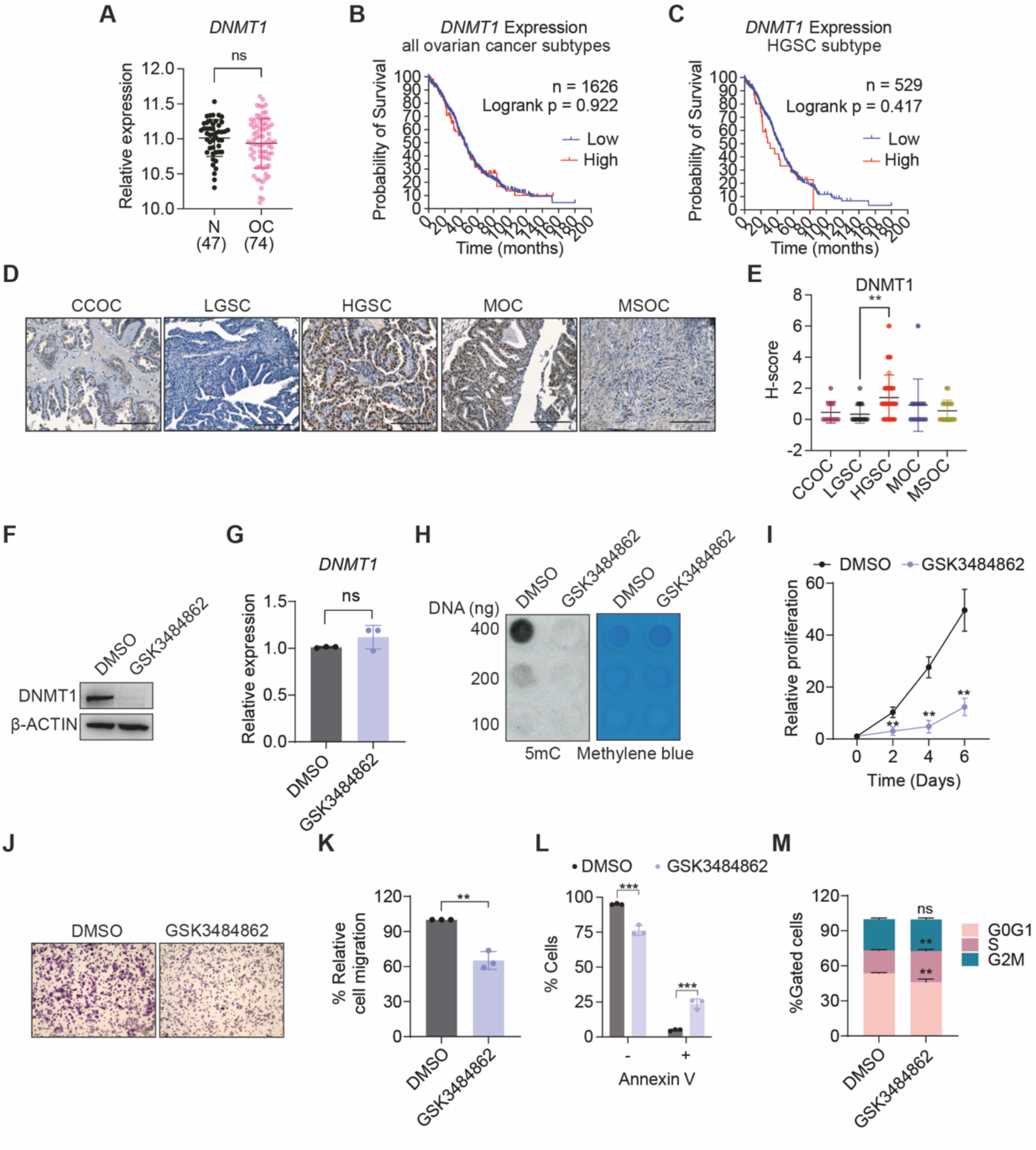
DNMT1 expression and functional impact in ovarian cancer. **A**. Scatter plot showing the expression levels of *DNMT1* in the dataset GSE37582. **B, C.** Kaplan-Meier plot illustrating the prognostic value of *DNMT1* expression in all ovarian cancer subtypes (**B**) and in the HGSC subtype (**C**), as determined using the TCGA database. **D**. Representative immunohistochemical staining images showing expression levels of DNMT1 across various ovarian cancer subtypes. CCOC-Clear Cell Ovarian Carcinoma; LGSC-Low Grade Serous Carcinoma; HGSC-High Grade Serous Carcinoma; MOC-Mucinous Ovarian Carcinoma; MSOC-Metastatic Ovarian Carcinoma. Scale bar, 50µM. **E**. Scatter plot depicting H-scores for DNMT1 immunohistochemical staining in ovarian cancer patient tissue samples analyzed in (**D**). **F, G.** Western blot image (**F**) and qPCR analysis (**G**) showing the expression levels of DNMT1 in GSK3484862-treated cells; β-ACTIN serves as a loading control. **H**. Representative dot blot demonstrating the 5mC levels in DMSO and GSK3484862-treated cells, with methylene blue staining used as a DNA loading control. **I**. Line graph depicting the effects of GSK3484862 on cell proliferation, illustrating the growth of DMSO and GSK3484862-treated cells over a six-day period. **J**. Representative trans-well assay images showing the migration of DMSO and GSK3484862-treated cells. Scale bar, 200 µm. **K**. Quantification of migrated cells from panel (**J**), calculated from absorbance values at 570 nm of dissolved crystal violet stain. **L**. Bar graph representing the percentage of apoptotic cells in DMSO and GSK3484862-treated cells, depicting the proportions of live (Annexin V-) and apoptotic (Annexin V+) cells. **M**. Cell cycle distribution of cells following GSK3484862 treatment. Data are presented as mean ± SD or as representative images from more than three independent biological experiments. Statistical significance was determined using Student’s t-test for comparisons between two groups, and one-way ANOVA for comparisons among three or more groups. ns = not significant (p > 0.05); *p < 0.05; **p < 0.01; ***p < 0.001.

Therefore, to further evaluate DNMT1 at the protein level, we performed immunohistochemical analysis on human tissue microarrays (TMAs) of 180 cases spanning various ovarian carcinoma subtypes (**Fig. 4D**). In contrast to the unchanged mRNA levels observed in bulk ovarian cancer tissues (**Fig. 4A**), DNMT1 protein expression was significantly elevated in HGSC compared to LGSC further reinforcing its contribution to HGSC pathogenesis (**Fig. 4E**).

To directly assess the functional role of DNMT1 in ovarian cancer, we treated OVCAR8 cells with GSK3484862, a selective DNMT1 inhibitor that promotes DNMT1 degradation via a proteasome-dependent mechanism without affecting its RNA levels (**Fig. 4F,G**) [21]. Dot blotting confirmed effective DNMT1 inhibition by GSK3484862, as evidenced by a significant reduction in 5-methylcytosine (5mC) levels (**Fig. 4H**). Phenotypically, GSK3484862 treatment reduced the growth and migration of OVCAR8 cells, induced apoptosis, and caused S-phase cell cycle arrest (**Fig. 4I-M**). Taken together, these findings underscore the oncogenic role of DNMT1 in ovarian cancer progression, particularly in HGSC.

### The METTL3–DNMT1 axis silences tumor suppressor genes in ovarian cancer

To identify DNMT1 target genes in ovarian cancer, we conducted RNA-seq on OVCAR8 cells following treatment with the DNMT1 inhibitor GSK3484862. This analysis identified 222 differentially expressed genes, including 196 up-regulated and 26 down-regulated genes in response to DNMT1 inhibition (**Fig. 5A, Table S4**). GO enrichment analysis of the upregulated genes showed significant enrichment in biological processes such as transcriptional regulation, apoptosis, chromatin remodelling, negative regulation of cell population proliferation and cellular response to hypoxia, among others (**Fig. 5B**).

**Figure 5.**
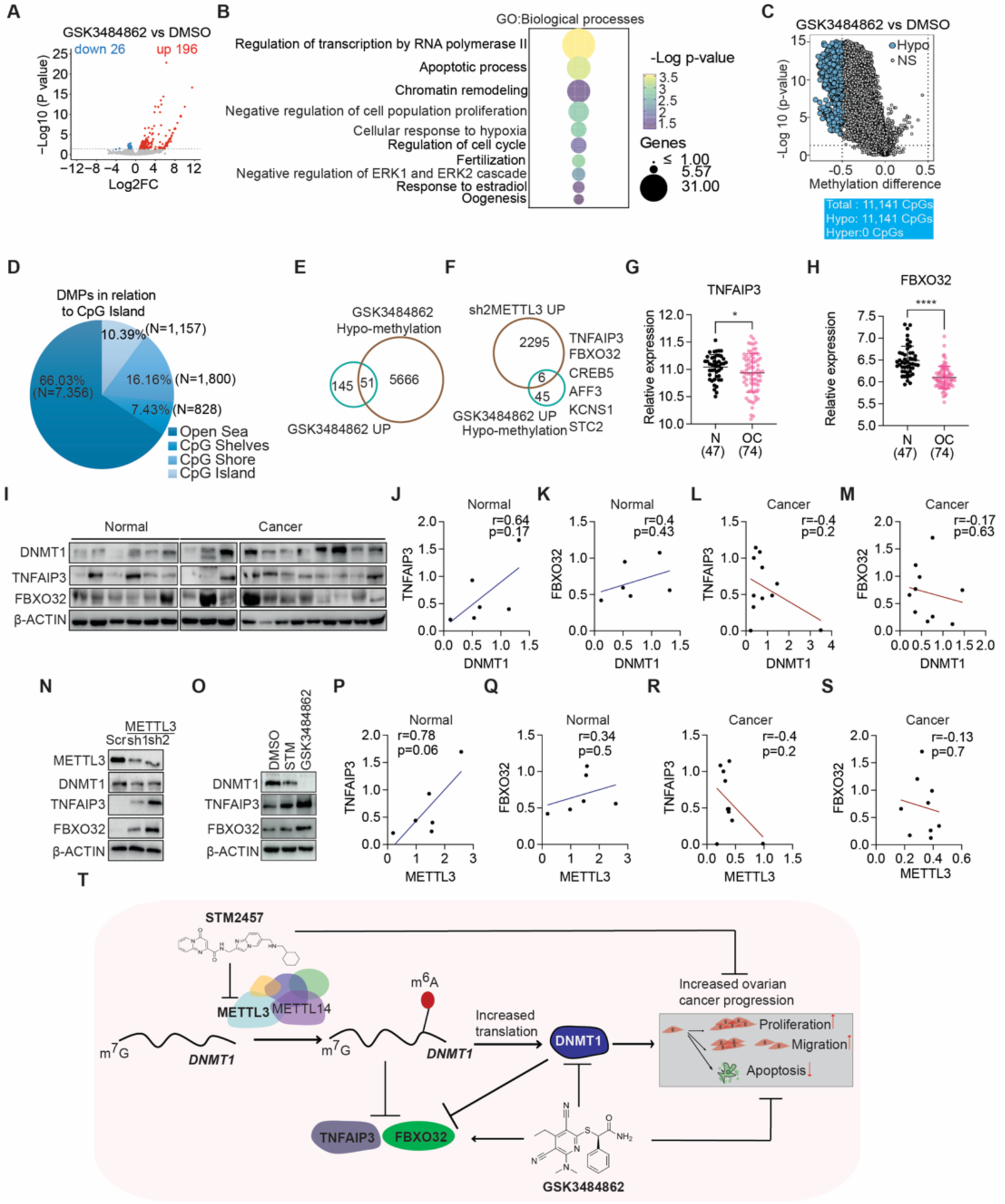
Epigenetic regulation of tumor suppressors by DNMT1. **A.** Volcano plot of differentially expressed genes in GSK3484862-treated cells compared to DMSO-treated OVCAR8 cells, highlighting significantly up-(red) and down-regulated (blue) genes. **B**. Dot plot showing enriched GO terms for genes that are up-regulated upon GSK3484862 treatment. **C**. Plot shows the methylation difference in GSK3484862 treated cells compared to DMSO control. Differentially methylated CpG (DMPs) sites were defined as those with an adjusted p-value ≤ 0.05 and a mean methylation difference of ≥ 0.5 between treatment conditions. **D**. Pie chart demonstrates the distribution of differentially methylated probes in relation to CpG island location. **E**. Venn diagram shows the overlap of upregulated genes with those associated with hypomethylated CpG sites following GSK3484862 treatment. **F**. Venn diagram depicting the overlap between genes that are upregulated and hypomethylated upon GSK3484862 treatment and those upregulated following METTL3 depletion (sh2). **G, H**. Scatter plots showing the expression levels of TNFAIP3 (**G**) and FBXO32 (**H**) in ovarian cancer patient samples compared to normal tissue, as analyzed using GEO dataset GSE37582. **I.** Western blot depicting DNMT1 expression and its target genes TNFAIP3 and FBXO32 in normal ovarian epithelial tissues and ovarian cancer tissues. β-ACTIN serves as a loading control. **J-M.** Plots show the correlation between DNMT1 protein expression and its target tumor suppressor genes in normal (**J-K**) and ovarian cancer tissues (**L-M**), based on quantification of protein levels from the western blot assay (**I**) using ImageJ. **N**. Immunoblot shows the expression of TNFAIP3 and FBXO32 in METTL3 depleted cells. β-ACTIN serves as a loading control. **O**. Expression of TNFAIP3 and FBXO32 following pharmacological inhibition of METTL3 and DNMT1. β-ACTIN serves as a loading control. **P-S**. Correlation plots demonstrate the correlation between the expression of METTL3 (corresponding to the expression observed in panel Fig. 3K) with TNFAIP3 and FBXO32 in normal (**P-Q**) and ovarian cancer tissues (**R-S**). **T**. Schematic diagram showing role of METTL3-DNMT1 axis in ovarian tumor progression. Data presented in as mean ± SD or as representative images from more than three independent biological experiments. Statistical significance was determined using Student’s t-test for comparisons between two groups. *p < 0.05; ****p< 0.0001.

To further explore the mechanisms underlying these transcriptional changes, we investigated whether DNMT1 inhibition leads to DNA hypomethylation at specific genomic loci by performing methylation array analysis on GSK3484862-treated OVCAR8 cells. t-distributed Stochastic Neighbor Embedding (t-SNE) analysis showed a clear separation between DMSO- and GSK3484862-treated samples (**Fig. S5A**). The DNA methylation B-value density plot displayed a bimodal distribution, with a higher peak for unmethylated CpGs, indicating a global reduction in DNA methylation following DNMT1 inhibition (**Fig. S5B)**. Consistently, the mean methylation levels were lower in GSK3484862-treated samples than in the controls (**Fig. S5C)**. Unsupervised analysis of all CpG sites in the array further confirmed global hypomethylation following treatment with the DNMT1 inhibitor (**Fig. S5D)**. 11,141 hypomethylated CpG sites were identified in response to GSK3484862 treatment (**Fig. 5C).** Examination of the genomic distribution of CpGs in relation to CpG islands revealed that most hypomethylated CpGs were found in open sea regions (66.03%), while smaller fractions were associated with CpG islands (10.39%), shelves (7.43%; 2–4 kb from the promoter CpG islands), and shores (16.16%; 0–2 kb from the promoter CpG islands) (**Fig. 5D**). In terms of gene location, 49.65% of differentially methylated CpG sites (DMPs) were found within gene bodies, while 25.2% were located in the 5′ regulatory regions, and 21.65% were in intergenic regions. The remaining 3.49% were situated in the 3′ untranslated regions (3′UTR) (**Fig. S5E**). When analyzed by gene type, 72.14% of the DMPs were located in coding regions, whereas 21.65% and 6.21% were found in intergenic and non-coding regions, respectively (**Fig. S5F**).

Integration of RNA-seq and methylation array data identified 51 (26%) overlapping transcripts, that were both upregulated and hypomethylated upon DNMT1 inhibition (**Fig. 5E**). To further investigate whether these transcripts were also regulated by METTL3, we focused on the more efficient knockdown achieved by sh2 METTL3 (**Fig. 5F).** Among the overlapping genes, we prioritized tumor necrosis factor alpha-induced protein 3 (TNFAIP3) and F-Box Protein 32 (FBXO32) because of their well characterized tumor suppressor roles [22–28].

To investigate the potential involvement of TNFAIP3 and FBXO32 in ovarian cancer, we first analyzed their expression patterns using the GEO dataset GSE37582. Both genes exhibited consistent downregulation in ovarian cancer tissue samples relative to controls, in line with their established roles as tumor suppressors (**Fig. 5G, H**). To explore their prognostic relevance, we conducted an overall survival analysis using TCGA data for patients with ovarian cancer. Although the observed associations did not reach statistical significance, there was a notable trend toward improved survival in patients with high TNFAIP3 and FBXO32 expression, particularly in the HGSC subtype (**Fig. S5G–J**).

Subsequent analysis of patient-derived ovarian tumor specimens revealed elevated DNMT1 expression, concomitant with reduced expression of TNFAIP3 and FBXO32, compared to normal ovarian tissues (**Fig. 5I**). Although not statistically significant, correlation analysis of normal ovarian samples demonstrated a positive association between DNMT1 expression and that of TNFAIP3 and FBXO32 (**Fig. 5J–K**). Conversely, this relationship was reversed in ovarian cancer tissues, in which DNMT1 expression was negatively correlated with the expression of both genes (**Fig. 5L–M**). These findings suggest that DNMT1-mediated epigenetic regulation contributes to the transcriptional repression of TNFAIP3 and FBXO32 in ovarian cancer.

Given the observed inverse correlation between DNMT1 and the tumor suppressors TNFAIP3 and FBXO32 in ovarian cancer, we next sought to validate the regulatory role of METTL3 in modulating their expression. Following METTL3 knockdown, western blot analysis demonstrated an increase in the protein levels of both TNFAIP3 and FBXO32, indicating that METTL3 negatively regulated their expression (**Fig. 5N**). Consistently, pharmacological inhibition of METTL3 using STM2457 elevated the expression of both genes to levels comparable to those observed with GSK3484862 (**Fig. 5O**). In line with the expression patterns associated with DNMT1, correlation analysis revealed a positive association between METTL3 and tumor suppressor genes in normal ovarian tissue, which shifted to a negative correlation in ovarian cancer samples (**Fig. 5P–S**) although these associations were not statistically significant. These parallel trends suggest a coordinated regulatory mechanism involving the METTL3–DNMT1 axis in the transcriptional repression of TNFAIP3 and FBXO32. Collectively, these findings support a model in which METTL3 and DNMT1 cooperatively mediate the epigenetic silencing of tumor suppressor genes, thereby contributing to ovarian tumor progression, particularly within the HGSC subtype (**Fig. 5T**).

## Discussion

RNA modifications, particularly m^6^A, are now recognized as critical regulators of gene expression. In the context of ovarian cancer, dysregulation of m^6^A “writers,” “readers,” and “erasers” contribute to tumorigenesis, metastasis, and chemoresistance [7–13, 29–40]. Our study builds on this model by revealing that METTL3 and METTL14 function as oncogenic drivers in HGSC, at least partially, through translational control of DNMT1.

Depletion of METTL3 or METTL14 impaired the proliferation and metastatic capacity of ovarian cancer cells both in vitro and in vivo, as evidenced by the reduced tumor growth in mice bearing ovarian cancer cells lacking either factor. Pharmacological inhibition of METTL3 by STM2457 recapitulated these effects in cells, underscoring its therapeutic potential. METTL3 and METTL14 form a heteromeric complex that mutually stabilizes their protein levels and functions as core m^6^A methyltransferases. Consequently, their canonical roles in m^6^A deposition are interdependent, complicating efforts to distinguish their individual contributions to HGSC. Nevertheless, transcriptomic analyses following individual knockdown revealed distinct gene expression signatures, suggesting potential m^6^A-independent, moonlighting functions. TCGA data further showed that METTL3, but not METTL14, was transcriptionally upregulated in ovarian cancer, indicating differential regulatory mechanisms.

m^6^A profiling revealed enrichment of m^6^A sites within CDS and near stop codons— consistent with previous studies [30], and implicated modified transcripts in pathways relevant to both ovarian development and cancer progression. Among these, DNMT1 has emerged as a key target, with m^6^A sites detected within its CDS. While prior work in non-small cell lung cancer showed increased DNMT1 expression with m^6^A methylation mediated by METTL3, this conclusion was based solely on correlative expression analyses [41]. In our study, we provide functional evidence that m^6^A enhances DNMT1 protein levels independently of mRNA abundance, supporting a translational regulatory mechanism. This was further supported by the decreased luciferase activity of a DNMT1 Exon 8 reporter following STM2457 treatment, and by diminished association of *DNMT1* mRNA with polysomes, suggesting impaired translational efficiency in the absence of m^6^A.

These findings align with recent reports that m^6^A modifications within CDS enhance translation in ovarian cancer [30], likely by resolving RNA secondary structures that impede elongation [42]. However, emerging evidence also links CDS-localized m^6^A to translation-coupled RNA decay under specific conditions, particularly when m^6^A residues occupy the ribosomal A site, causing ribosomal stalling and subsequent transcript relocation to P-bodies [43]. Thus, the outcome of m^6^A modification is context-dependent and must be evaluated for each specific transcript.

While Quarto et al., 2025 showed that the METTL3-METTL14 complex recruits DNMT1 to chromatin to mediate 5mC deposition in gene bodies in HeLa cells and mouse embryonic stem cells [44], our study uncovered a distinct regulatory mechanism in HGSC. Here, METTL3 and METTL14 regulate DNMT1 post-transcriptionally through m^6^A-dependent enhancement of translation. Nonetheless, we cannot exclude the possibility that the chromatin-associated roles of METTL3/14 also contribute to DNMT1 regulation in this cancer context.

Given the central role of DNMT1 in ovarian cancer [45, 46], our findings linking METTL3/METTL14-mediated m^6^A modification to DNMT1 translation have important therapeutic implications. DNA methyltransferase inhibitors (DNMTis), including FDA-approved agents such as 5-azacytidine and 5-aza-2′-deoxycytidine, inhibit DNMT activity and promote protein degradation, with proven efficacy in hematologic malignancies. In this study, we demonstrated that the novel DNMT1 inhibitor GSK3484862 exerts anti-tumor activity in HGSC.

Our data suggest that the METTL3/METTL14–DNMT1 axis functions as a regulatory node repressing tumor suppressor genes, such as TNFAIP3 and FBXO32, sustaining oncogenic gene expression programs in HGSC. Disrupting this axis—either by inhibiting METTL3 to suppress DNMT1 translation or by using DNMTis to degrade DNMT1—may impair tumor growth and reverse epigenetic silencing of tumor suppressors. Together, these findings reveal a key epitranscriptomic mechanism by which METTL3 enhances the translation of epigenetic regulators such as DNMT1 and highlight the therapeutic potential of combining epigenetic and epitranscriptomic strategies in ovarian cancer.

## Material and Methods

### Datasets

To examine *METTL3*, *METTL14*, *DNMT1*, *TNFAIP3* and *FBXO32* expression in patient samples, the GSE37582 dataset from the GEO database was analyzed by comparing the mRNA expression profiles of 74 ovarian cancer cases with 47 healthy controls. Additionally, TCGA datasets were used to assess the correlation between *METTL3*, *METTL14*, *DNMT1*, *TNFAIP3,* and *FBXO32* expression levels and overall survival in patients with ovarian cancer.

### Cell culture and treatment

OVCAR8 cells were grown in Roswell Park Memorial Institute (RPMI) 1640 medium supplemented with 10% fetal bovine serum (FBS) and 1% penicillin/ streptomycin (Gibco). HEK-293T cells, used for lentiviral production, were cultured in Dulbecco’s Modified Eagle Medium (DMEM) (Gibco) supplemented with 10% FBS and 1% penicillin/streptomycin. All the cells were maintained at 37°C and 5% CO_2_ in a humidified incubator. For METTL3 inhibition, cells were treated with STM2457 (MedChemExpress; HY-134836, 10µM) for 48 h. To assess RNA stability, cells were treated with actinomycin D (5 µg/ml) for specified times, followed by RNA extraction.

### Antibodies

For western blot analysis, the following antibodies were used at their respective dilutions: Anti-METTL3 (Abcam, ab221795, 1:1000); Anti-METLL14 (Sigma, HPA038002, 1:1000), Anti-KIAA1429 (Abcam, ab50993, 1:1000), Anti-CBLL1 (Abcam, ab50993, 1:2000), Anti-WTAP (Invitrogen, OTI12G, 1:2000), Anti-DNMT1 (Abcam, ab188453, 1:1000), Anti-UHRF1 (ThermoFisher Scientific, PAF-29884, 1:1000). The secondary antibodies included goat Anti-Rabbit IgG H&L (HRP) (Abcam, ab6721, 1:5000) and goat Anti-Mouse IgG H&L (HRP) (Abcam, 1:1000, ab6789). Anti-β-actin (Sigma, A5441, 1:2000) or β-tubulin (Sigma, T2200, 1:2000) were used as loading controls for western blotting. For IF staining, the following antibodies were used: Anti-METTL3 (Abcam, ab221795, 1:250); Anti-METLL14 (Sigma, HPA038002, 1:250), Anti-KIAA1429 (Abcam, ab50993, 1:100), Anti-CBLL1 (Abcam, ab50993, 1:250), Anti-WTAP (Invitrogen; OTI12G, 1:100), Anti-DNMT1 (Abcam; ab188453, 1:250), Anti-5mC (Active Motif; 39649, 1:250). The secondary antibodies used for IF included goat anti-rabbit IgG AF568 (ThermoFisher Scientific, A11011, 1:1000), and goat anti-mouse IgG AF488 (ThermoFisher Scientific, A11029, 1:1000).

### Human ovarian cancer specimens and immunohistochemistry

To investigate the correlation between the expression of METTL3, DNMT1, TNFAIP3, and FBXO32 in patient samples, 11 ovarian cancer tissues along with six normal ovarian tissues were used in the study (**Supplementary Table S5**). For protein extraction, tissues were finely minced and suspended in 300 μl lysis buffer containing 50 mM HEPES (pH 7.5), 150 mM NaCl, 3 mM MgCl₂, 0.2% Triton X-100, 0.2% Nonidet P-40, and 10% glycerol, supplemented with a protease inhibitor cocktail. The tissues were homogenized using a tissue homogenizer and centrifuged at 12000 rpm for 15 minutes. The resulting supernatant containing total protein was quantified using standard Bradford assay and used for immunoblotting.

Immunohistochemical analysis was performed on ovarian cancer tissue microarrays (TMAs) obtained from Biomax (OV1006 and OV802c819) using a primary antibody against DNMT1 (Abcam, ab188453). The OV1006 TMA consists of 100 cores representing 100 individual cases, including 62 serous carcinomas, 10 mucinous adenocarcinomas, five clear cell carcinomas, three endometrioid adenocarcinomas, ten lymph node metastases, and ten adjacent normal ovarian tissue samples. The OV802c819 TMA comprised 40 serous carcinomas, six clear cell carcinomas, two mucinous adenocarcinomas, one stromal tumor, two sex cord-stromal tumors, nine germ cell tumors, ten metastatic adenocarcinomas, and ten adjacent normal ovarian tissue samples. Although both TMAs included adjacent normal ovarian tissues, epithelial cells were not present in these sections; therefore, the normal samples were excluded from the final analysis. DNMT1 expression was quantified using the H-score, calculated as the product of staining intensity and percentage of positively stained cells. The staining intensity was graded on a scale from 0 to 3 (0 = no staining, 1 = weak, 2 = moderate, 3 = strong), while the percentage of positive cells was scored from 0 to 4 (0 = 0%, 1 = 1–25%, 2 = 26–50%, 3 = 51–75%, 4 = 76–100%).

### Generation of *METTL3* and *METTL14* KD OVCAR8 cells

Lentiviral particles were produced by transfecting HEK-293T cells with the packaging vector pCMVdR8.2, the helper plasmid pCMV-VSV-G and pLKO.1-puro plasmids containing shRNAs targeting *METTL3*, *METTL14* or a *Scramble* (Scr) control sequence (target sequences provided in **Supplementary Table S6**). Transfections were carried out using jetPEI (Polyplus) according to the manufacturer’s protocol or CaCl_2_. Lentiviral particles containing the supernatant were harvested 48 and 72 h post-transfection. After 48 h of supernatant collection, the cells were supplemented with fresh medium. Lentiviral particles were concentrated using Amicon Ultra-15 Centrifugal Filter Units (Merck, Darmstadt, Germany). OVCAR8 cells were transduced with *Scr*, *METTL3* and *METTL14* shRNA containing lentiviral particles in the presence of polybrene (8 ug/ ml). The transduced cells were selected with puromycin (0.1 ug/ml) for 2 days post-infection.

### Cell proliferation assay

2 × 10⁴ cells, depleted of METTL3 or METTL14 and treated with either STM2457 or GSK3484862 as indicated, were seeded into six-well plates. The cell numbers were quantified on days 2, 4, and 6 after seeding. Proliferation rates were calculated and normalized to Scr, and relative growth curves were plotted.

### Colony forming assay

Cells depleted of METTL3 or METTL14 and treated with STM2457 or GSK3484862 as indicated, were seeded at low density (1 × 10³ cells per well) in six-well plates for colony formation assays. The cells were cultured for 10 days, and the medium was replenished every 48 h. The colonies were fixed with 10% formaldehyde and stained with 0.25% crystal violet. Plates were digitally imaged, the crystal violet-stained colonies were solubilized in 10% acetic acid for quantification. The absorbance was measured at 570 nm using a microplate reader (Tecan, Thermo). Colony growth was expressed as the percentage of cell viability relative to Scr.

### Trans-well migration assay

Cells depleted of METTL3 or METTL14, and treated with STM2457 or GSK3484862 as indicated, were cultured in serum-free growth medium for 48 h to induce cell cycle arrest and minimize proliferation. Subsequently, 5 × 10⁵ serum-starved cells were seeded into the upper chambers of 12-well transwell inserts (Corning), whereas the lower chambers were filled with growth medium containing 10% FBS to serve as a chemoattractant. After 24 h of migration, the cells on both sides of the membrane were fixed with 10% formaldehyde and stained with 0.25% crystal violet. Non-migrating cells on the upper surface of the membrane were removed using a sterile cotton swab. The migrated cells on the lower membrane surface were visualized using a Leica microscope at 20× magnification. For quantification, migrated cells were counted in six random fields per membrane using the ImageJ Cell Counter plugin. Data are plotted as the average number of migrated cells relative to Scr.

### Cell Cycle and apoptosis assay

Cell cycle and apoptosis analyses were conducted using the BD Accuri C6 Plus Flow Cytometer and the Muse Cell Analyzer (Millipore, Sigma-Aldrich) respectively, following the manufacturer’s protocols. For cell cycle analysis using BD Accuri C6 Plus, the cells were stained with propidium iodide (PI; ThermoFisher). For apoptosis assays, Annexin V Ready Flow Conjugates for Apoptosis Detection (ThermoFisher) were used in combination with eBioscience™ 7-AAD Viability Staining Solution, according to the manufacturer’s instructions.

### qRT-PCR

Total RNA was extracted using the RNeasy Plus Mini Kit (Qiagen) or TRIzol reagent, as previously described [47]. Equal amounts of RNA were used for complementary DNA (cDNA) synthesis using the RevertAid First Strand cDNA Synthesis Kit (Invitrogen), following the manufacturer’s protocol. Quantitative real-time PCR (qRT-PCR) was performed using gene-specific primers (**Supplementary Table S6**) and SYBR Green Master Mix (ThermoFisher Scientific) on the QuantStudio Real-Time PCR System (Applied Biosystems). β-actin served as an internal control for normalization. Cycle threshold (Ct) values were analyzed using the 2^–ΔΔCt method to calculate the relative fold changes across at least three independent biological replicates.

### Immunoblotting

Whole-cell lysates were prepared by incubating the cells in lysis buffer containing 50 mM HEPES (pH 7.5), 150 mM NaCl, 3 mM MgCl₂, 0.2% Triton X-100, 0.2% Nonidet P-40, and 10% glycerol, supplemented with a protease inhibitor cocktail. Proteins were resolved on 8–10% SDS-PAGE gels and transferred to PVDF membranes (0.45 µm, Merck Millipore). Membranes were blocked with 5% skim milk (Invitrogen) for 1 h at room temperature, followed by overnight incubation with primary antibodies at 4°C. After washing, membranes were incubated with horseradish peroxidase-conjugated secondary antibodies for 1 h. Protein bands were visualized using enhanced chemiluminescence (Thermo Scientific) and imaged using an Amersham™ Imager 680.

### Immunofluorescence

METTL3 depleted cells were seeded onto poly-L-lysine–coated coverslips in 12-well plates, fixed with 4% formaldehyde (Sigma-Aldrich), and permeabilized using 0.25% Triton X-100 (Sigma-Aldrich). Following two washes with PBS, the cells were blocked for 1 h at room temperature in blocking solution containing 10% goat serum (Invitrogen), 1% BSA (Hyclone), and 0.01% Tween-20 (Sigma-Aldrich). Subsequently, cells were incubated with METTL3 and METTL14 primary antibodies overnight at 4°C. After washing with PBS, cells were incubated with fluorescence-conjugated secondary antibodies. Nuclei were counterstained with 4′,6-diamidino-2-phenylindole (DAPI). Fluorescent images were acquired using a Leica SP8 confocal microscope and analyzed using the ImageJ software.

### RNA sequencing and differential gene expression analysis

RNA sequencing for METTL3 and METTL14 knockdown was performed using Novogene - Advancing Genomics, Improving Life (https://en.novogene.com/) using two biological replicates. RNA sequencing analysis of GSK3484862 treated OVCAR8 cells was conducted by Biomarker Technologies (BMKGENE) (www.bmkgene.com) with three biological replicates. Differentially expressed genes (DEGs) were identified from the combined dataset using a log_2_ fold-change threshold (upregulation >0.58; downregulation <−0.58) and a significance cut-off of p-adj <0.05 (−log_10_(p-adj) >1.3).

### Gene Ontology (GO) Analysis

GO analysis of DEGs was performed using the web-based tool Database for Annotation, Visualization, and Integrated Discovery (DAVID).

### Xenograft study

Luciferase-labelled OVCAR8 cells (1 × 10^6^) were subcutaneously injected into 6-7-week-old female Crl:NU(NCr)-Foxn1^nu^ nude mice. Images at day 0 were taken immediately after cell injection. In vivo bioluminescence imaging was performed using a PerkinElmer IVIS system (model INVCLS136335) with Living Image software (v4.3.1). For imaging, luciferin (150 ug/mL, 100ul) was retro-orbitally injected into the mice. Tumor volume was measured with a digital caliper using the following formula: tumor volume = 1/2 x (length × width^2^). Tumors were collected at the experimental endpoint for further analysis.

### MeRIP sequencing

MeRIP sequencing was performed as previously described [48]. Briefly, fragmented mRNA (5 µg) was immunoprecipitated overnight with m^6^A antibody (10 µg, Abcam, ab151230). Protein A Dynabeads® (61006, Thermo Fisher Scientific) was used to capture the m^6^A modified mRNA. The beads were then washed and the pool of m^6^A modified transcripts was eluted in 16 µl of DEPC-treated water. 4 µl of the eluate was used to prepare cDNA using the SuperScript™ VILO™ cDNA Synthesis Kit (Thermo Fisher), and qRT-PCR was performed for *GAPDH* and *MYC* to validate the immunoprecipitation. 10 µl (∼100 ng) of the eluate were sent to Novogene (https://en.novogene.com/) for library preparation and sequencing. Integrated Genome Browse (IGV) was used to study the m^6^A peaks of specific mRNAs.

### SELECT

Site-specific m^6^A detection of the DNMT1 transcript was performed using a single-base elongation- and ligation-based qPCR amplification method (SELECT), as previously described [20]. Up- and down- probes were designed based on the MeRIP-seq coordinates of m^6^A and m^6^A-6 sites (as input) on DNMT1. Briefly, mRNA was purified from total RNA extracted from DMSO-, STM2457-treated, METTL3-and METTL14-depleted OVCAR8 cells using Protein A Dynabeads® (61006, Thermo Fisher Scientific). The mRNA was first annealed with the respective up- and down- probes using dNTPs (N0447S, NEB) and Cutsmart buffer (B6004S, NEB) by incubating at the following decreasing temperature gradient: 90°C 1 min, 80°C 1 min, 70°C 1 min, 60°C 1 min, 50°C 1 min and 40°C 6 min in a thermocycler (Applied Biosystems). The m^6^A-dependent gap filling and ligation was facilitated by incubating annealed samples with Bst 2.0 DNA Polymerase (M0537S, NEB), SplintR ligase (M0375S, NEB) and Adenosine 5’-Triphosphate (ATP) (P0756S, NEB) in a thermocycler for 20 min at 40°C followed by 20 min at 80°C. SELECT products were quantified by qRT-PCR using primers targeting a small overhang sequence present on each up/ down probes. ΔCT values were calculated by normalizing the CT value for each m^6^A site to its input. ΔΔCT values were obtained by normalizing STM2457- treated samples to the ΔCT of DMSO controls. The relative SELECT products were calculated by inverting the log transformation using the 2-^ΔΔCt^. The sequences of the probes and primers are listed in **Supplementary Table S6.**

### Polysome profiling

METTL3-depleted and Scr control cells were treated with cycloheximide (CHX, 100 µg/ml) for 5 min and washed twice with cold 1x PBS containing CHX to halt translation. Equal numbers of cells were lysed in polysome lysis buffer (20 mM Tris-HCl, pH 7.4, 5 mM MgCl2, 150 mM NaCl, 1% Triton X-100) supplemented with CHX (100 µg/ml), DTT (1 mM), protease inhibitors (complete EDTA-free, Roche), and RNase Inhibitor (200 U/ml, Themo Scientific). Lysates were agitated on a rocking platform for 10 min and then centrifuged at 10,000 g for 10 min at 4°C. Equal volumes of clarified lysates were layered onto sucrose gradients (15-45% w/v) that were previously prepared using a Gradient Master (BioComp Instruments). The lysates were centrifuged at 40,000 g for 2 h at 4°C in an ultracentrifuge with an SW60Ti rotor (Beckman). Fractions were collected using a Gilson Fraction FC-203B collector in Eppendorf tubes containing 300 µl of solution II (10 mM Tris-HCl pH 7.5; 350 mM NaCl; 10 mM EDTA; 1% SDS; 42% urea), while measuring the absorbance at 260 nm using a Triax flow cell (BioComp Instruments). Polysome profiles were generated by plotting absorbance against piston position.

### Luciferase reporter assay

The DNMT1 Exon 8 sequence was amplified by PCR (Phusion™ High-Fidelity DNA Polymerase, Thermo Scientific™) from cDNA obtained from the OVCAR8 cell line using the RevertAid kit (Thermo Fisher) with oligo(dT) primers. The cDNA was then digested using XhoI and NotI and inserted into the multiple cloning site of the psiCheck2 plasmid, which was digested with the same restriction enzymes and purified using the QIAquick Gel Extraction Kit (Qiagen). The cloned DNMT1 Exon 8-psiCheck2 construct was validated using Sanger sequencing. The primers and sequences used are listed in **Supplementary Table S6**. For the luciferase assay, OVCAR8 cells were seeded in 12-well plates and transfected with DNMT1 Exon 8-psiCheck2 reporter plasmid or empty vector using Lipofectamine LTX according to the the manufacturer’s instructions (Thermo Fisher). Cells were treated with the METTL3 inhibitor STM2457 or DMSO (control) 24 h post-transfection. The dual luciferase assay was carried out 48 h after STM2457 treatment using the Dual-Luciferase® Reporter Assay System (Promega) according to the manufacturer’s instructions. Data were normalized to the value of Renilla divided by firefly luciferase. Luciferase activity was calculated in STM2457-treated cells relative to DMSO-treated cells.

### MethylationEPIC v2.0 BeadChip hybridization

DNA bisulfite conversion and array hybridization were conducted at the Genomics Unit of Josep Carreras Leukemia Research Institute (Badalona, Spain). Total DNA was quantified using the Qubit dsDNA Broad Range Assay Kit (Invitrogen™, Thermo Fisher Scientific, MA, USA), according to the manufacturer’s instructions. Subsequently, 600 ng of DNA was bisulfite-converted using the EZ-96 DNA Methylation Kit (Zymo Research Corp., CA, USA) following the manufacturer’s protocol. Finally, 200 ng of bisulfite-converted DNA was hybridized onto the Infinium HumanMethylationEPIC version 2.0 BeadChip microarray (Illumina Inc.), as previously described in [49].

### DNA methylation analysis

All analyses were conducted using the R statistical software (R version 4.1.2) (www.R-project.org), following the minfi pipeline [50]. Briefly, raw signal intensity files (IDATs) were processed, and sample quality was evaluated by excluding samples with a mean detection p-value ≥ 0.05. Next, normalization was performed using the Noob method, and low-quality probes were filtered out. Specifically, probes with a mean detection p-value ≥ 0.01 in more than 10% of samples, cross-reactive probes, as well as those associated with sex chromosomes and SNPs, were removed from the analysis. Probes with missing values were imputed based on the mean methylation values of the corresponding probes from other samples. Finally, methylation beta-values and M-values were generated for downstream analysis. Unsupervised hierarchical clustering was performed using all CpG probes to cluster the samples using the Euclidean distance and the ward. D2 clustering method. For row clustering, a random subset comprising 1% CpG probes was selected to reduce computational complexity. A heatmap was generated and visualized using the ComplexHeatmap package in R [51]. t-distributed Stochastic Neighbor Embedding (t-SNE) analysis was performed on all CpG sites in the array using the Rtsne R package [4]. Global methylation analysis was performed by calculating the mean methylation value for each sample, which was then plotted and grouped by condition using the ggplot2 R package [52].

To identify differentially methylated probes, a linear regression model was applied to methylation M-values and one condition was compared. CpG sites were considered statistically significant if the adjusted p-value was ≤ 0.05. Additionally, a biological filter was applied, considering only probes that were statistically significant and a mean methylation difference of at least |0.5| between the conditions. The annotation of differentially methylated probes was performed using the UCSC annotation (GRCh38/hg38). Gene set enrichment analysis was performed using the EnrichR R package [6] for genes associated with differentially methylated CpG sites identified in the previous analysis. The analysis was performed using Gene Ontology databases including “Biological Process 2023,” “Molecular Function 2023,” and “Cellular Component 2023.” A gene set was considered significant if the adjusted p-value was ≤ 0.05.

### Statistical analysis

All data are shown as ±SD (Standard Deviation). Graphs were plotted and statistical analyses were performed using GraphPad Prism (version 10). Student’s *t*-test and one-way ANOVA were used to calculate significance. \**p*< 0.05, \*\**p* < 0.01, \*\*\**p*< 0.001 \*\*\*\**p* <0.0001 were considered as statistically significant.

## Ethics statement

All ovarian cancer specimens were collected from the biobank of Norrlands Universitetssjukhus, Umeå, Sweden. All samples were handled in accordance with the approved biobank protocol (Dnr: 2019-03729). The Ethics Committee for Animal Experimentation (CEEA-PRBB) had approved all animal experiments conducted at the Hospital del Mar Medical Research Institute, Barcelona, Spain.

## Acknowledgements

We would like to thank Aguilo Lab members for their support and cooperation, the Biochemical Imaging Center Umeå and the National Microscopy Infrastructure, NMI (VR-RFI 2019-00217) for assistance with the imaging processing. We thank S. Gildlund who performed IHC staining.

## Funding

This research was supported by grants from the Knut and Alice Wallenberg Foundation, Umeå University, Västerbotten County Council, Swedish Research Council (2017-01636; 2022-01322), Kempe Foundation (SMK-1766, JCK-1723.1 and JCK-2150), Cancerfonden (190337 Pj; 222455 Pj), and Cancer Research Foundation in Northern Sweden (AMP 21-1030).

## Author contributions

KK and FA conceived and designed the study. KK performed most of the experiments. IPN and TCT conducted the in vivo (mice) experiments. AN and ME performed DNA methylation analyses. CP, ED, ACR and AN performed the bioinformatics analysis. GK conducted the protein stability assays. KP and CH performed mass spectrometry analyses. AP and EL evaluated the immunohistochemical staining of the tissue microarrays. KK and FA wrote the original draft of the manuscript. All authors contributed to the data interpretation and reviewed and edited the final manuscript.

## Data availability

All next-generation sequencing data can be publicly accessed in ArrayExpress or GEO webserver. Methylation data were submitted to the GEO webserver with ID GSE299247.

## Conflict of interest statement

None declared.

## Notes

### Competing Interest Statement

The authors have declared no competing interest.

