## Supplementary figures S1-S5 and Table S5-S6 for "METTL3-mediated m^6^A modification of DNMT1 enhances ovarian cancer progression"

### Index of Supplemental Figures:

- **Figure S1:** Knockdown efficiency of METTL3 and METTL14 and the effect of pharmacological inhibition of METTL3 in OVCAR8.
- **Figure S2:** Impact of METTL3 depletion on methyltransferase complex integrity and tumor burden, and the prognostic significance of elevated METTL3 and METTL14 in ovarian cancer.
- **Figure S3:** Differentially expressed genes upon METTL3 and METTL14 depletion in OVCAR8 cells.
- **Figure S4:** Impact of METTL3 and METTL14 depletion on DNMT1 expression
- **Figure S5:** GSK3484862 treatment leads to hypomethylation of DNA in OVCAR8 cells and elevated TNFAIP3 and FBXO32 levels are associated with improved patient overall survival in ovarian cancer

### Figure S1

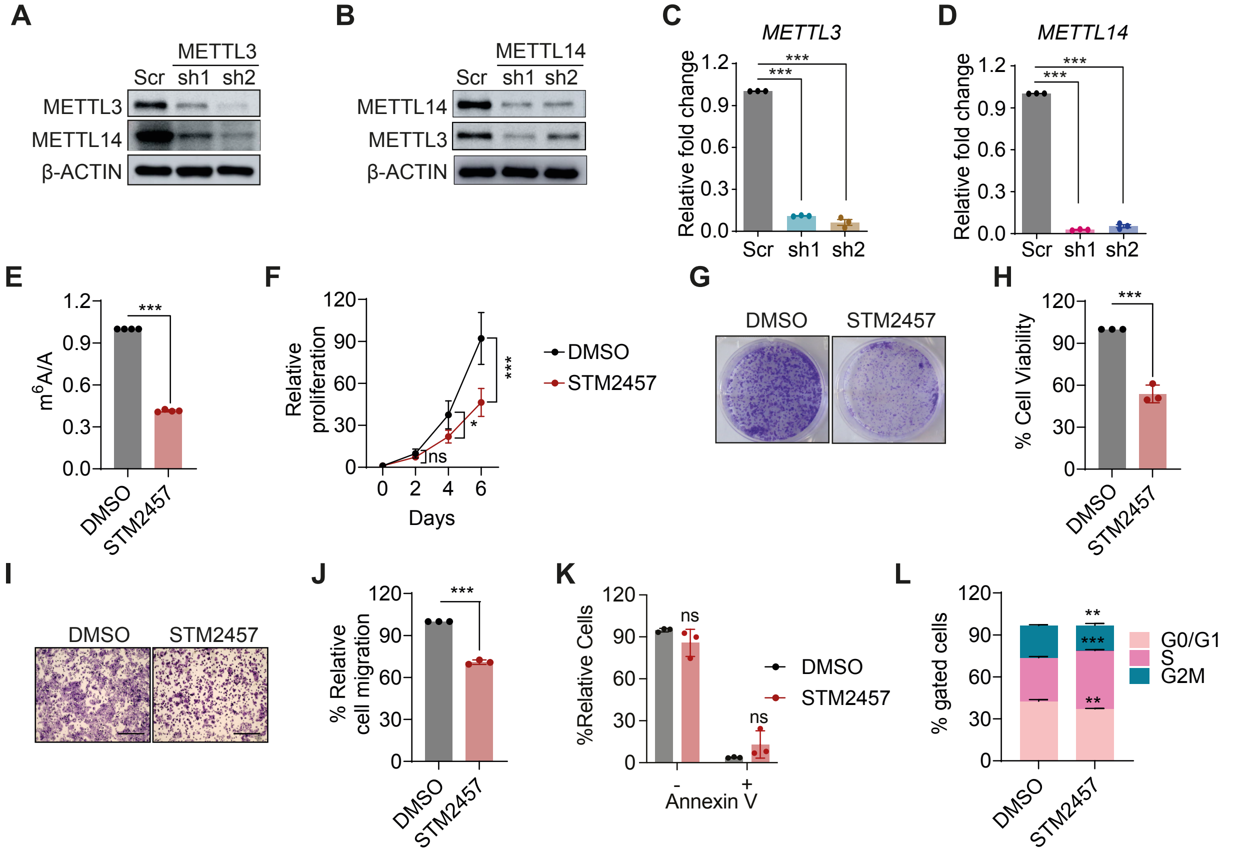


**Figure S1: Knockdown efficiency of METTL3 and METTL14 and the effect of pharmacological inhibition of METTL3 in OVCAR8. A**, **B**. Representative Western blot images showing knockdown efficiency of METTL3 (**A**) and METTL14 (**B**). β-ACTIN was taken as loading control. **C, D**. Bar graphs showing qPCR analysis of *METTL3* (**C**) and *METTL14* (**D**) mRNA knockdown efficiency. β-actin was used as a loading control. **E**. Levels of $m^6A$ upon STM2457 treatment as measured by quantitative LC-MS/MS. **F**. Line graph depicts the growth of DMSO and STM2457-treated cells over a 6-day period. **G**. Representative images from the colony formation assay conducted over 10 days, demonstrating the ability of OVCAR8 cells to form colonies in the presence of DMSO or STM2457. **H**. Bar graph depicting relative percent cell viability based on absorbance values at 570 nm from dissolved crystal violet, corresponding to the colony formation assay in panel (**G**). **I**. Representative trans-well assay images illustrating the migration of cells treated with DMSO or STM2457. Scale bar, 200 µm. **J**. Quantification of migrated cells from panel (**I**), calculated from absorbance values at 570 nm of dissolved crystal violet stain. **K**. Bar graph displaying the percentage of apoptotic cells in DMSO- and STM2457-treated OVCAR8 cells, illustrating the proportion of live (Annexin V–) and apoptotic (Annexin V+) cells. **L**. Cell cycle distribution of cells following STM2457 treatment is demonstrated by the bar graph. Data are presented as mean ± SD or as representative images from more than three independent biological experiments. Statistical significance was determined using Student's t-test for comparisons between two groups, and one-way ANOVA for comparisons among three or more groups. ns = not significant ($p > 0.05$); *$p < 0.05$; **$p < 0.01$; ***$p < 0.001$.

**Figure S2**

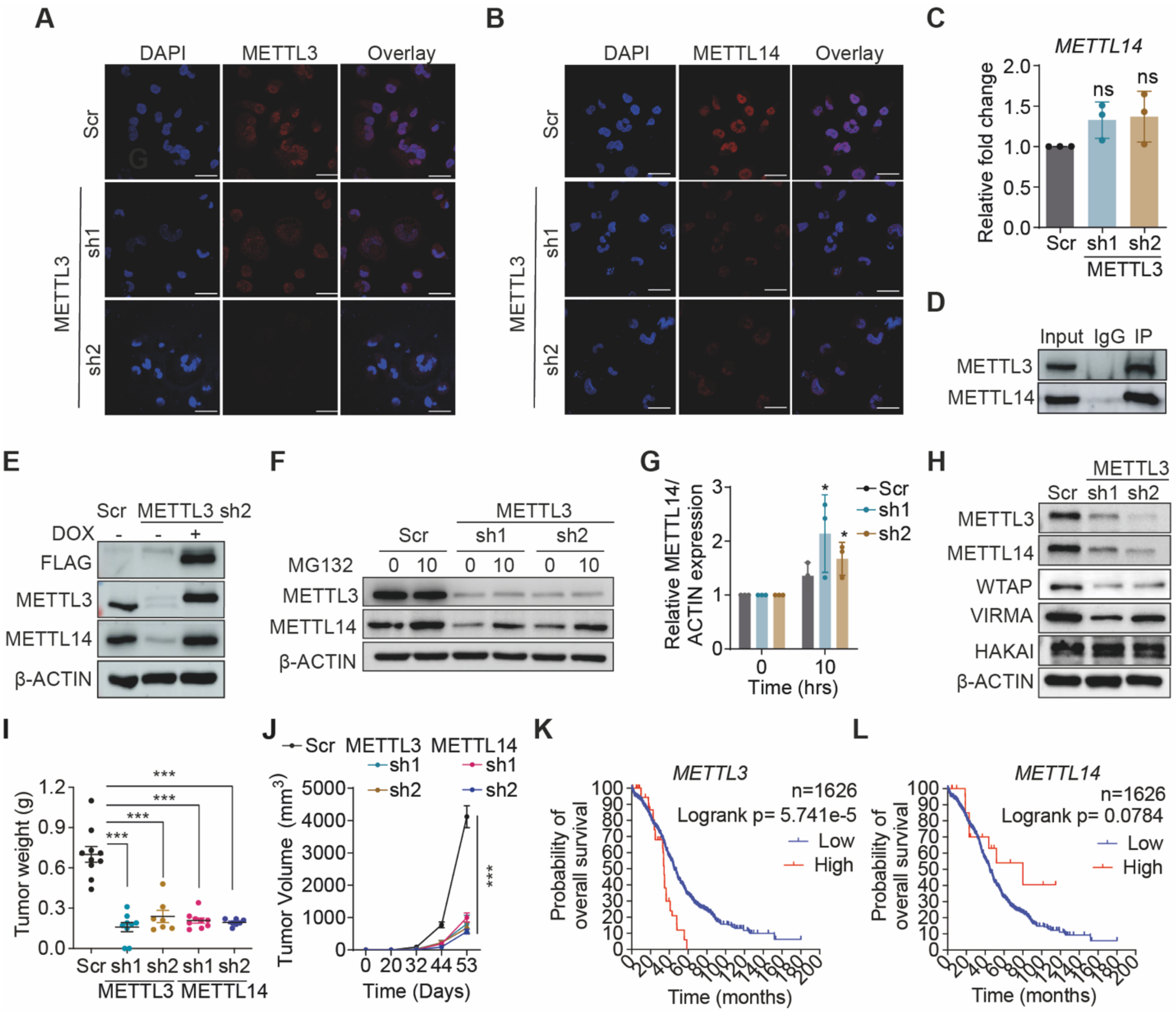


**Figure S2: Impact of METTL3 depletion on methyltransferase complex integrity and tumor burden, and the prognostic significance of elevated METTL3 and METTL14 in ovarian cancer. A**, **B**. Immunofluorescence images depicting the expression of METTL3 and METTL14 in METTL3-depleted cells. **C**. qPCR analysis of *METTL14* following METTL3 knockdown. β-actin was used as a loading control. **D**. Immunoblot showing the interaction between METTL14 and METTL3 following immunoprecipitation with a METTL3 antibody. Rabbit IgG was used as a negative control. **E**. Western blot of FLAG, METTL3 and METTL14 from whole-cell extracts from scr and METTL3-knockdown (sh2) cells and upon expression of wild-type cytoplasmic METTL3 (sh2 + doxycycline (dox)) cells. **F**. Immunoblot showing the expression of METTL3 and METTL14 in METTL3-depleted cells following treatment with MG132 for 10 hours; β-ACTIN is shown as the loading control. **G**. Graph shows the normalized protein expression levels of METTL14 observed in (**F**). **H**. Western blot images showing expression levels of METTL3, METTL14, WTAP, VIRMA, and HAKAI upon METTL3 knockdown; β-ACTIN was used as a loading control. **I**. Graph displays weight of tumors collected from mice transplanted with control, METTL3- and METTL14-depleted OVCAR8 cells corresponding to the tumors shown in panel (**Fig.1O**). **J** Line graph displays the growth of Scr control and METTL3- and METTL14-depleted OVCAR8 tumors in BALB/c nude mice at the indicated time points during the study period. **K**, **L**. Kaplan-Meier survival curves for overall survival for METTL3 (**K**) and METTL14 (**L**) expression in all subtypes of ovarian cancer using TCGA data. Data are presented as mean ± SD or as representative images from more than three independent biological experiments. Statistical significance was determined using Student's t-test for comparisons between two groups, and one-way ANOVA for comparisons among three or more groups. ns = not significant ($p > 0.05$); *$p < 0.05$; **$p < 0.01$; ***$p < 0.001$.

**Figure S3**

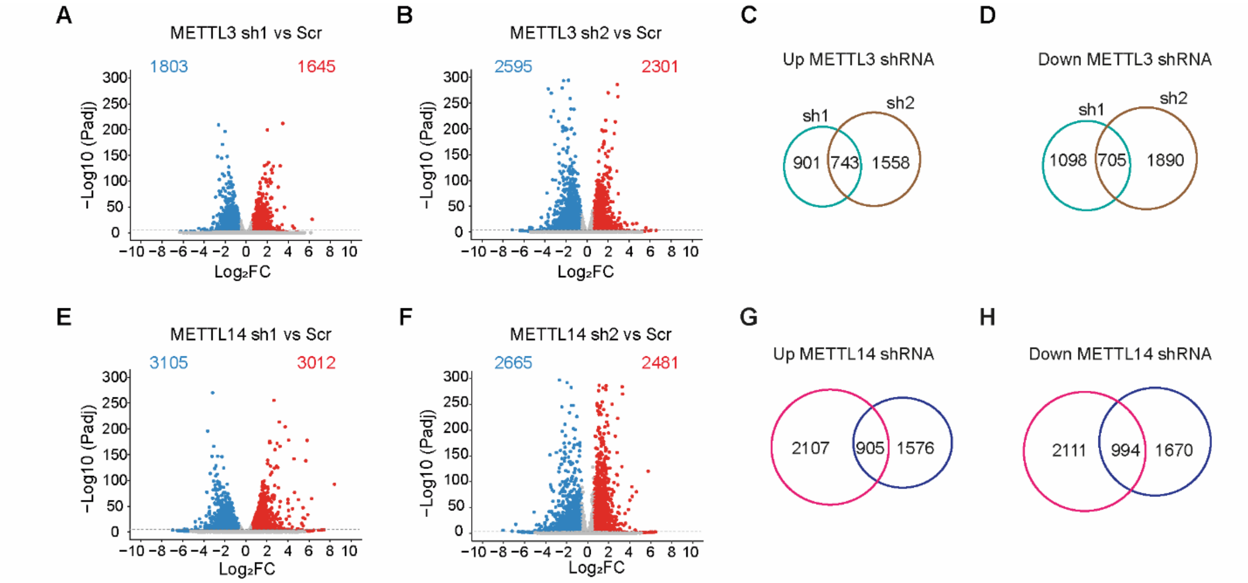


**Figure S3: Differentially expressed genes upon METTL3 and METTL14 depletion. A, B**. Volcano plots of differentially expressed genes in sh1METTL3- (**A**) and sh2METTL3-depleted (**B**) OVCAR8 cells compared to Scr cells, respectively, highlighting significantly up- and down-regulated genes. **C, D** Venn diagram showing commonly upregulated (**C**), and downregulated (**D**) genes in METTL3 OVCAR8 cells. **E, F** Volcano plots of differentially expressed genes in sh1METTL14- (**E**) and sh2METTL14-depleted (**F**) OVCAR8 cells compared to Scr cells, respectively, highlighting significantly up- and down-regulated genes. **G, H** Venn diagram showing commonly upregulated (**G**) and downregulated (**H**) genes in METTL14 OVCAR8 cells. Representative images of n=2 independent biological experiments.

**Figure S4**

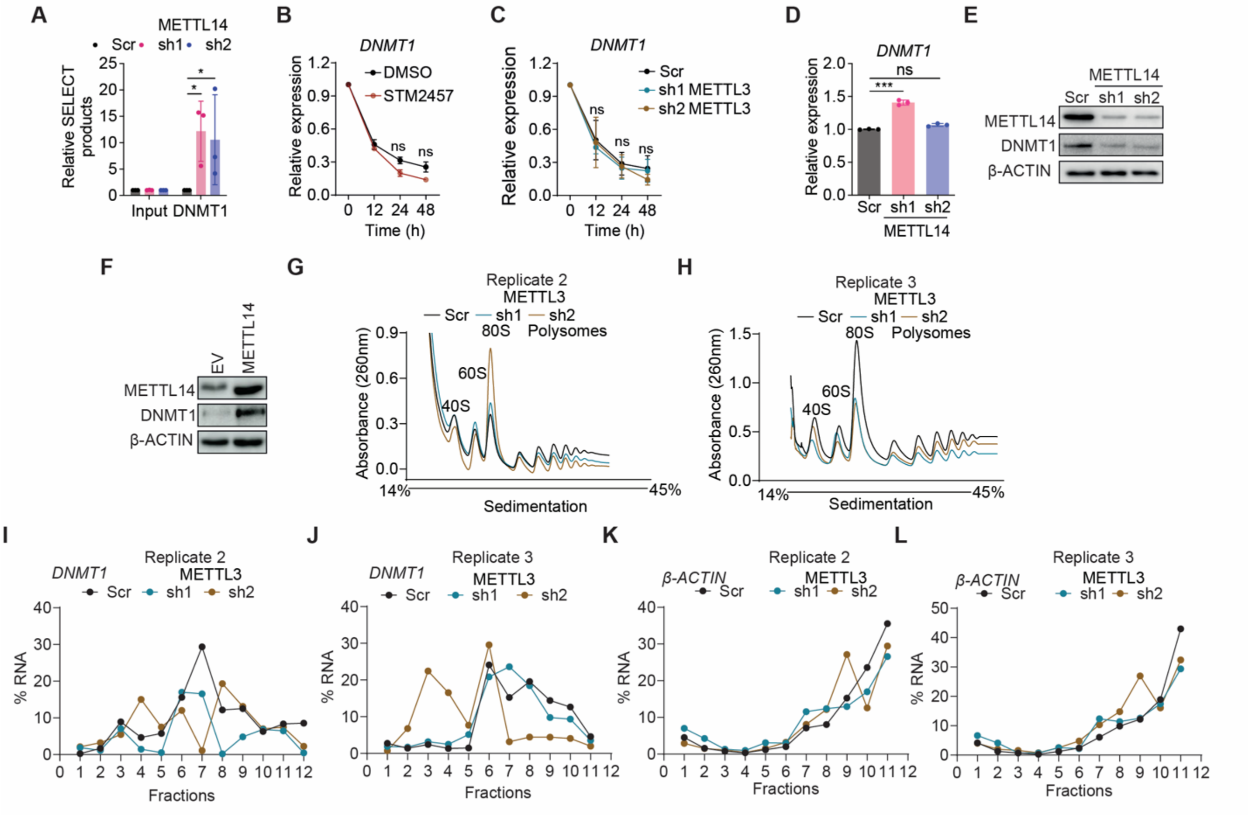


**Figure S4: Impact of METTL3 and METTL14 depletion on DNMT1 expression. A.** Bar graphs illustrating the levels of SELECT products upon METTL14 depletion**. B, C.** Line graphs depicting the effects of Actinomycin D treatment upon STM2457 exposure (**B**) and METTL3 depletion (**C**), observed at 12-, 24- and 48-hours after treatment. **D.** qPCR analysis of *DNMT1* expression upon METTL14 depletion. **E.** Immunoblot showing DNMT1 protein levels in METTL14-depleted cells. β-ACTIN was used as a loading control. **E**. Western blot image depicts the expression level of DNMT1 upon overexpression of METTL14. **G, H.** Polysome profiling curve for METTL3-depleted cells, illustrating global ribosomal distribution. **I-L.** qPCR analysis of *DNMT1* (**I-J**) and *β-ACTIN* (**K-L**) mRNA in polysome fractions following profiling shown in (**G-H**). Data are presented as mean ± SD or as representative images from more than three independent biological experiments. Statistical significance was determined using Student's t-test for comparisons between two groups, and one-way ANOVA for comparisons among three or more groups. ns = not significant ($p > 0.05$); *$p < 0.05$.

**Figure S5**

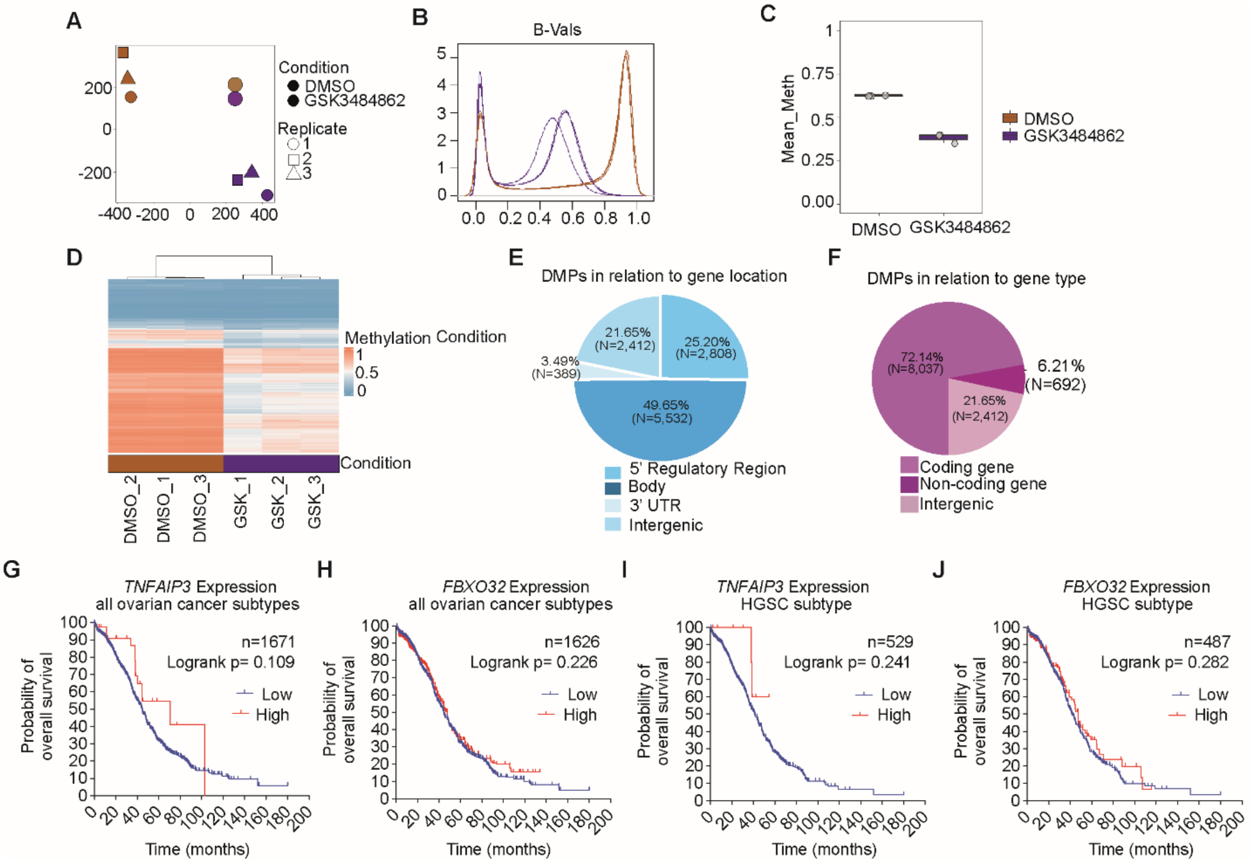


**Figure S5. GSK3484862 treatment leads to hypomethylation of DNA in OVCAR8 cells and elevated TNFAIP3 and FBXO32 levels are associated with improved patient overall survival in ovarian cancer.** **A**. t-distributed Stochastic Neighbor Embedding (t-SNE) analysis plot depicting the clustering of DMSO and GSK3484862-treated experimental groups in three biological replicates. **B.** Beta density plot for samples with orange curves representing DMSO control and violet curves representing GSK3484862 treated group. **C**. Scatter plot shows the mean-methylation levels in DMSO and GSK3484862-treated OVCAR8 cells. **D**. Heatmap displays the unsupervised hierarchical analyses of the global DNA methylome of the DMSO and GSK3484862-treated groups. **E, F.** Pie chart demonstrates the distribution of differentially methylated probes in relation to gene location (**E**) and gene type (**F**). **G, H.** Overall survival analysis of *TNFAIP3* and *FBXO32* across all ovarian cancer subtypes using data from the TCGA database. **I, J**. Overall survival analysis of *TNFAIP3* and *FBXO32* in HGSC subtype of ovarian cancer. Methylation data are presented as representative images of n=3 independent biological experiments

**Index of Supplemental Tables:**

**Supplementary Table**

- **Table S1:** RNA-seq analysis data for METTL3 KD cells compared to Scr control. Refer to the separate Excel file, Table S1.xlsx
- **Table S2**: RNA-seq analysis data for METTL14 KD cells compared to Scr control. Refer to the separate Excel file, Table S2.xlsx
- **Table S3**: MeRIP analysis data for OVCAR8 wild type. Refer to the separate Excel file, Table S3.xlsx
- **Table S4**: RNA-seq analysis data for GSK3484862 treated OVCAR8 cells compared to DMSO control. Refer to the separate Excel file, Table S4.xlsx
- **Table S5**: Details of ovarian patient samples used in the study.
- **Table S6:** Primers, constructs and shRNAs sequences used in this study

**Table S5.** Details of ovarian patient samples used in the study.

| **Patient sample details** | | |
|---|---|---|
| **Patient id** | **Tissue type/ Histology subtype** | **Histology grade** |
| 224b | Normal | NA |
| 234d | Normal | NA |
| 239d | Normal | NA |
| 242d | Normal | NA |
| 252d | Normal | NA |
| 254d | Normal | NA |
| 209a | Ovarian carcinoma, HGSC | 3 |
| 212a | Ovarian carcinoma, HGSC | 3 |
| 220s | Ovarian carcinoma, HGSC | 3 |

| | | |
|---|---|---|
| 221a | Ovarian carcinoma, HGSC | 3 |
| 226a | Ovarian carcinoma, HGSC | 3 |
| 240a | Ovarian carcinoma, HGSC | 3 |
| 245a | Ovarian carcinoma, HGSC | 3 |
| 248a | Ovarian carcinoma, HGSC | 3 |
| 243a | Ovarian carcinoma, HGSC | 3 |
| 252a | Ovarian carcinoma, HGSC | 3 |
| 254a | Ovarian carcinoma, HGSC | 3 |

**Table S6.** Primers, constructs and shRNAs sequences used in this study

| **shRNAs** | | |
|---|---|---|
| **Target Gene** | **Plasmid** | **Sequence 5´→ 3´** |
| Scramble | pLKO.1-puro-shScramble | CAACAAGATGAAGAGCACCAA |
| *METTL3* | pLKO.1-puro-sh*METTL3*_1 | GCAAGTATGTTCACTATGAAA |
| | pLKO.1-puro-sh*METTL3*_2 | CGTCAGTATCTTGGGCAAGTT |
| *METTl14* | pLKO.1-puro-sh*METTL14*_1 | GCCGTGGACGAGAAAGAAATA |
| | pLKO.1-puro-sh*METTL14*_2 | GCTAATGTTGACATTGACTTA |

| **qPCR** | | |
|---|---|---|
| **Gene** | **Forward primer 5´→ 3´** | **Reverse primer 5´→ 3´** |
| *Mettl3* | AACTGCAACGCATCATTCGG | TTGACACCAACCAAGCAGTG |
| *METTL14* | AGGGGTTGGACCTTGGAAGA | GAAGTCCCCGTCTGTGCTAC |
| *β-actin* | ACCAACTGGGACGACATGGAG AAG | TACGACCAGAGGCATACAGGG ACA |
| *DNMT1* | GTGGCCCAGACTAGGAGGCG | CCGTGATGGTCCGGAAAGGA C |

| **Primers for SELECT of DNMT1 m$^6$A site** | |
|---|---|
| *m$^6$A site_up* | tagccagtaccgtagtgcgtgTCCACGTTCCCCACCCCCTG |
| *m$^6$A site_down* | CCCCACGTCCTGGAAGGAAAcagaggctgagtcgctgcat |
| *Input_up* | TagccagtaccgtagtgcgtgTTCCCCACCCCCTGTCCCCA |
| *Input_down* | GTCCTGGAAGGAAATTGGGAcagaggctgagtcgctgcat |
| *qPCR_F for SELECT* | ATGCAGCGACTCAGCCTCTG |

| | |
|---|---|
| *qPCR_R for SELECT* | TAGCCAGTACCGTAGTGCGTG |

| **Luciferase reporter assay** | |
|---|---|
| psiCHECK2 | All predicted $m^6A$ sites in *Renilla* and *Firefly Luciferase* were mutated. |
| XhoI DNMT1_Exon8_F | GTCGAGTCAGGGCTCCTGAGATAATGA |
| NotI-DNMT1_Exon8_R | GCGGCCGCTTTCGATCACTAGCAAAACCCA |
| Sequencing primer | GCCTAAGATGTTCATCGAGTC |
